# Dissecting the role of CAR signaling architectures on T cell activation and persistence using pooled screening and single-cell sequencing

**DOI:** 10.1101/2024.02.26.582129

**Authors:** Rocío Castellanos-Rueda, Kai-Ling K. Wang, Juliette L. Forster, Alice Driessen, Jessica A. Frank, María Rodríguez Martínez, Sai T. Reddy

## Abstract

Chimeric antigen receptor (CAR) T cells represent a promising approach for cancer treatment, yet challenges remain such as limited efficacy due to a lack of T cell persistence. Given its critical role in promoting and modulating T cell responses, it is crucial to understand how alterations in the CAR signaling architecture influence T cell function. Here, we designed a combinatorial CAR signaling domain library and performed repeated antigen stimulation assays, pooled screening and single-cell sequencing to investigate T-cell responses triggered by different CAR architectures. Parallel comparisons of CAR variants, at early, middle and late timepoints during chronic antigen stimulation systematically assessed the impact of modifying signaling domains on T cell activation and persistence. Our data reveal the predominant influence of membrane-proximal domains in driving T cell phenotype. Additionally, we highlight the critical role of CD40 costimulation in promoting potent and persistent T cell responses, followed by CTLA4, which induces a long-term cytotoxic phenotype. This work deepens the understanding of CAR T cell biology and may be used to guide the future engineering of CAR T cell therapies.

## INTRODUCTION

Chimeric antigen receptor (CAR) T cells are an emerging therapeutic strategy for cancer treatment. CARs are synthetic receptors consisting of an extracellular antigen binding domain fused to intracellular signaling domains that trigger and modulate T cell responses upon activation. The infusion of genetically engineered CAR T cells in patients guides the recognition of a target tumor antigen and promotes tumor clearance while inducing long-lasting memory immunity (*1*, *2*). To date, six CAR T cell therapies have been approved by the FDA for the treatment of hematological malignancies, and there are over a thousand ongoing clinical trials for a broad range of cancer types (*3*). Despite the potential of these therapies, they still face several challenges, including associated toxicities, poor tumor infiltration, exhaustion and lack of T cell persistence, which have limited their clinical success in many indications (*4*).

In recent years, the search for solutions has motivated the engineering of different CAR designs that enable novel recognition and activation properties (*5*). In particular, the essential role of intracellular signaling elements in orchestrating cellular responses and the large diversity of existing immune signaling proteins have been harnessed to expand the repertoire of CAR signaling architectures. The architecture can be defined as the choice, number and specific arrangement (membrane proximal or distal) of signaling elements within the CAR construct. Moving beyond clinically approved CARs, which combine the signaling domains of the CD3ζ chain of the T cell receptor (TCR)/CD3 co-receptor complex and costimulatory receptors CD28 or 4-1BB, several studies have investigated the impact of making precise changes in the choice, number and order of signaling domains (*6–10*) or motifs (*11*, *12*). Despite technical limitations of functional assays, which restrict the number of constructs that can be individually produced and tested, pre-clinical studies have identified new CAR designs with distinct antitumor properties. For instance, combining CD79A and CD40 signaling domains resulted in CARs exhibiting improved proliferation and superior in vivo antitumor activity compared to clinically approved designs (*6*). Furthermore, incorporation of CTLA4 cytoplasmic tails into a CD28-CD3ζ CAR increased its cytotoxic potential while delaying T cell activation and proinflammatory cytokine production, ultimately enhancing CAR-T efficacy in a murine model of leukemia (*7*).

To further explore the vast CAR signaling domain combination space, several recent studies have designed high-throughput screening approaches to engineer CARs with distinct or enhanced functional properties. These strategies combine the use of signaling domain libraries, pooled screening, deep or single-cell sequencing and computational tools to address challenges in CAR T cell engineering. The choice of the optimal methodology, however, poses a non-trivial task. Employing different library designs and T cell platforms (primary cells or cell lines), Goodman et al. and Gordon et al. conducted pooled phenotypic screens through fluorescence-activated cell sorting (FACS) and deep sequencing to assess the enrichment of functional variants (*13*, *14*). Daniels et al. performed arrayed screening on a subset of a CAR library, recording flow cytometry-based phenotypic information, which was followed by machine learning to predict the cytotoxicity and memory potential of a larger library of signaling architectures (*15*). Notably, our group has performed pooled functional screening of a large CAR signaling domain library and used single-cell RNA sequencing (scRNAseq) for high-throughput assessment of T cell transcriptional phenotypes (*16*).

Currently, there is still limited understanding of how the architecture of a CAR translates to the functional or transcriptional phenotype of T cells. In addition, the dynamics of how such cellular phenotypes evolve over time requires further investigation, especially in a clinically-relevant context such as chronic antigen stimulation, which is known to drive T cell exhaustion, a main cause of therapy failure (*17*, *18*). Here, we systematically study the role of CAR signaling architectures on T cell activation and persistence by combining pooled functional screening of a combinatorial signaling domain library with scRNAseq. This enables the characterization of CAR T cell responses in a high-throughput manner, while mimicking the early and late stages of chronic tumor stimulation through an in vitro model of CAR T cell dysfunction. Capturing different single-cell transcriptomic snapshots across time, our data reveal intriguing patterns, such as the prominent influence of domains proximal to the cell membrane in modulating T cell phenotype and the pivotal role of CD40 costimulation in driving a potent yet persistent T cell response. Thus our study synergizes signaling domain engineering, pooled functional screening and scRNAseq to enhance the mechanistic understanding of CAR T cell signaling.

## RESULTS

### Design of a combinatorial signaling domain library of CAR variants

In order to systematically investigate the impact of modifying the intracellular architecture of CARs on T cell function, we generated a combinatorial signaling domain library based on 1st, 2nd or 3rd generation CAR designs; a classification based on the number of costimulatory domains (Fig. 1a). All CAR designs possessed the same extracellular domain consisting of a single-chain variable fragment (scFv) with binding specificity for the human epidermal growth factor receptor 2 (HER2), which is a tumor-associated antigen present on several solid cancers (*19*). The CD3ζ activation domain was combined with costimulatory signaling domains of five different immune receptors, which cover different receptor families that are known to trigger distinct signaling pathways for modulating T cell activity. CD28 and 4-1BB (tumor necrosis factor receptor superfamily 9; TNFRSF9) were selected as they are the most commonly used costimulatory domains and are present in clinically approved CAR T cell therapies. In addition, we included the signaling domains of CD40 (TNFRSF5) and the cytokine receptor chain IL15RA, which in preclinical studies have demonstrated the ability to enhance the anti-tumor properties of CARs (*6*, *20*, *21*). Lastly, CTLA4 was chosen as an example of an inhibitory receptor on T cells that still may enhance anti-tumor responses when incorporated into CARs (*7*). As a negative control, a non-signaling CAR (NS-CAR) was designed that lacks any intracellular signaling domains and is therefore unable to initiate CAR-dependent T cell activation. This results in a library with 32 different designs: one 1st generation, five 2nd generation, 25 3rd generation CARs and the NS-CAR as negative control (Fig. 1a).

**Figure 1:**
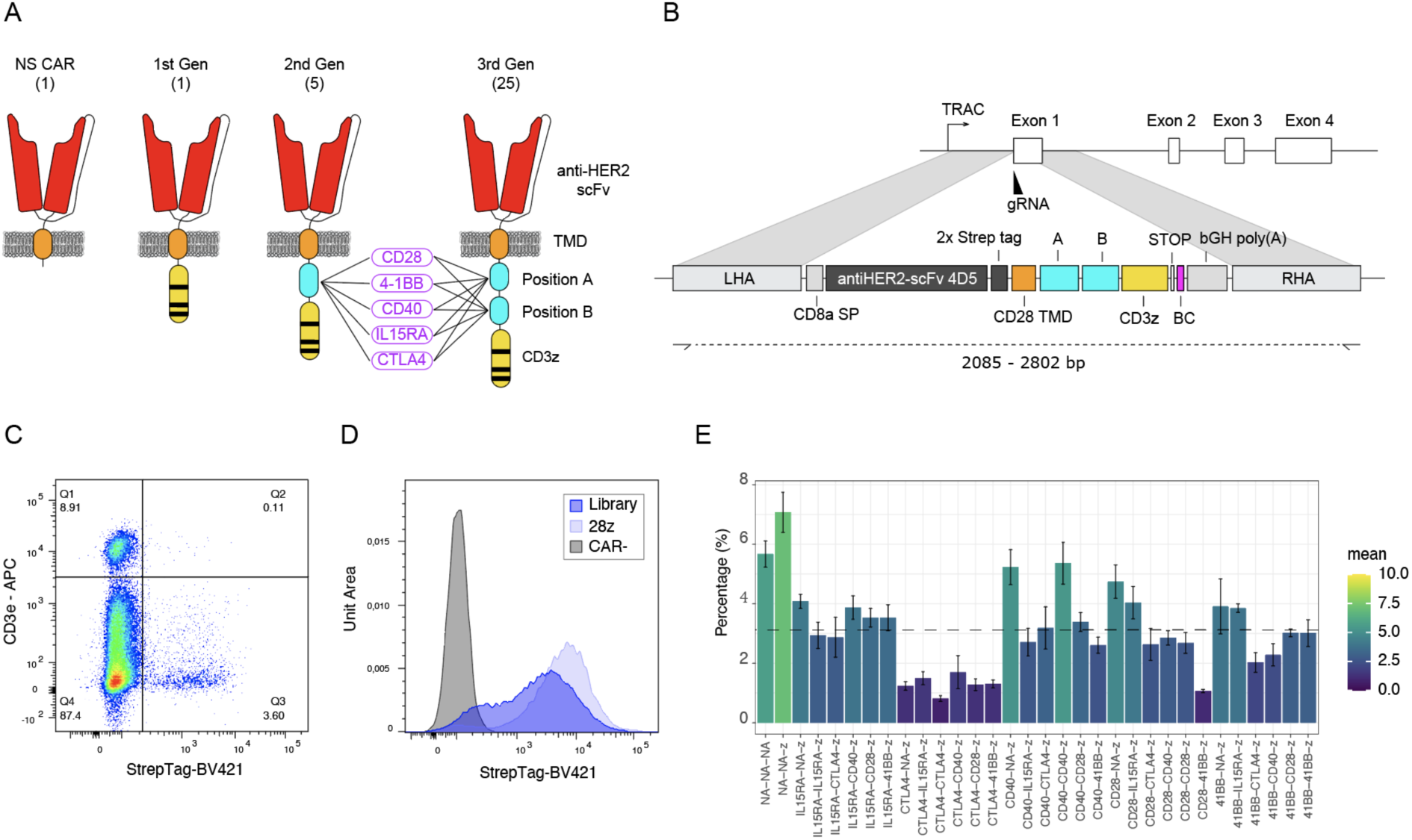
Design and production of a combinatorial signaling domain library of CAR variants. **A**- Schematic representation of the CAR library design. The library consists of 2nd and 3rd generation CAR designs that incorporate five selected costimulatory domains, which are shuffled in all possible combinations. The library also includes a 1st generation CAR and a non-signaling (NS) CAR that lacks signaling domains. When referring to domain positioning within the CAR, positions A and B denote domains located proximal or distal to the cell membrane, respectively. All variants contain an anti-HER2 single- chain variable fragment (scFv) and a CD28 transmembrane domain (TMD). **B**- Schematic shows the targeted genomic integration of the CAR library into the *TRAC* locus of T cells. Following a CRISPR- Cas9 guide RNA (gRNA)-directed double-stranded break at the start of exon 1 of the *TRAC* locus, a dsDNA repair template possessing left and right homology arms (LHA, RHA) and a full CAR gene (signal peptide (SP), scFv, TMD, signaling domains and poly-A signal) is used to induce homology-directed repair (HDR) and CAR gene insertion. **C**- Flow cytometry plot illustrating the T cell product obtained six days after the engineering of the CAR library into primary human T cells. Positive surface expression of a CAR (StrepTag) and negative expression of the TCR (CD3ε co-receptor) identifies correctly engineered CAR T cells. **D**- Flow cytometry histograms display CAR surface expression profiles of 28z and the pooled library of CAR T cells after enrichment compared to unedited T cells. **E**- Library diversity of the CAR T cell final product following enrichment and a 12-day expansion, assessed by deep sequencing. The dashed line represents the theoretically balanced distribution of the library. Barplot shows the mean of five biological replicates (CAR T cell products engineered from different healthy donors) and error bars represent SEM.

Next, we used CRISPR-Cas9 and homology-directed repair (HDR) to genomically integrate the CAR library into the TCR alpha chain (*TRAC*) locus of primary human T cells (Fig. 1b). Precise integration of the CAR gene into the *TRAC* locus ensures that every variant is expressed under the same transcriptional regulation while simultaneously knocking out the TCR (*22*), an appropriate setting to compare library candidates in a pooled manner. Following genome editing, engineered T cells were selected based on positive surface expression of a CAR (StrepTag) and negative expression of the TCR (CD3ε co-receptor) using FACS (Fig. 1c). To verify the quality of the engineered CAR-T cell product and validity of the library controls, we first examined the CAR surface expression and cytotoxic potential of T cells engineered with the CD28 2nd generation CAR (28z) or the negative control NS-CAR. Both CAR-T cell products displayed similar levels of CAR surface expression after enrichment (Sup. Fig 1a). Subsequently, T cell killing potential was measured by monitoring the growth curves of SKBR3 cells, a HER2-positive breast cancer cell line, following a 48h co-culture. As expected, 28z CAR T cells were able to efficiently eliminate all tumor cells, while NS- CAR T cells were unresponsive (Supp. Fig. 1b).

We next proceeded to produce a pooled library of CAR T cells including all 32 CAR variants. Surface expression of the CAR library in sorted T cells appeared to be more heterogeneous compared to the 28z CAR (Fig. 1d), indicating CAR variant-specific differences in cell surface expression. This is in particular expected for CARs containing a CTLA4 domain, where the presence of an endocytosis motif has been previously described to drive receptor recycling and degradation (*23*). Targeted deep sequencing of the CAR library confirmed that all variants were expressed and could be enriched by FACS. Except for a few variants that showed a lower enrichment, most of which indeed contained the signaling domain of CTLA4, the library variants were distributed at similar levels (Fig. 1e; CAR nomenclature is described in Supp. Table 1).

### Assessment of library persistence following repeated antigen stimulation

Next, we characterized CAR signaling domain variants using in vitro repeated antigen stimulation (RAS), an experimental workflow that aims to mimic chronic antigen stimulation from tumor cells (*24*, *25*), which is associated with CAR T cell exhaustion during clinical treatment. The pooled library of CAR T cells was repeatedly challenged with HER2-expressing SKBR3 cells for 12 days. Every third day, a sample of the co-cultured cells was restimulated with fresh SKBR3 cells and their effector potential was assessed by flow cytometry based on surface expression of the degranulation marker CD107a and intracellular expression of pro-inflammatory cytokines IFNγ and TNFα (Fig. 2a). At an early stage of the RAS assay (day 3), the CAR T cell library showed robust effector potential as evidenced by high degranulation and cytokine production (Fig. 2b). A consistent and gradual decline of this effector phenotype was observed towards later time points, indicating that the RAS assay could effectively recapitulate the progressive exhaustion of CAR T cells. Throughout the assay, CD8 T cells seemed to lose effector potency faster than CD4 T cells. In line with this observation, the fraction of CD8 T cells consistently dropped in time in favor of CD4 T cells, which seemed to have a longer lifespan in the context of an in vitro RAS co-culture (Fig. 2c).

**Figure 2:**
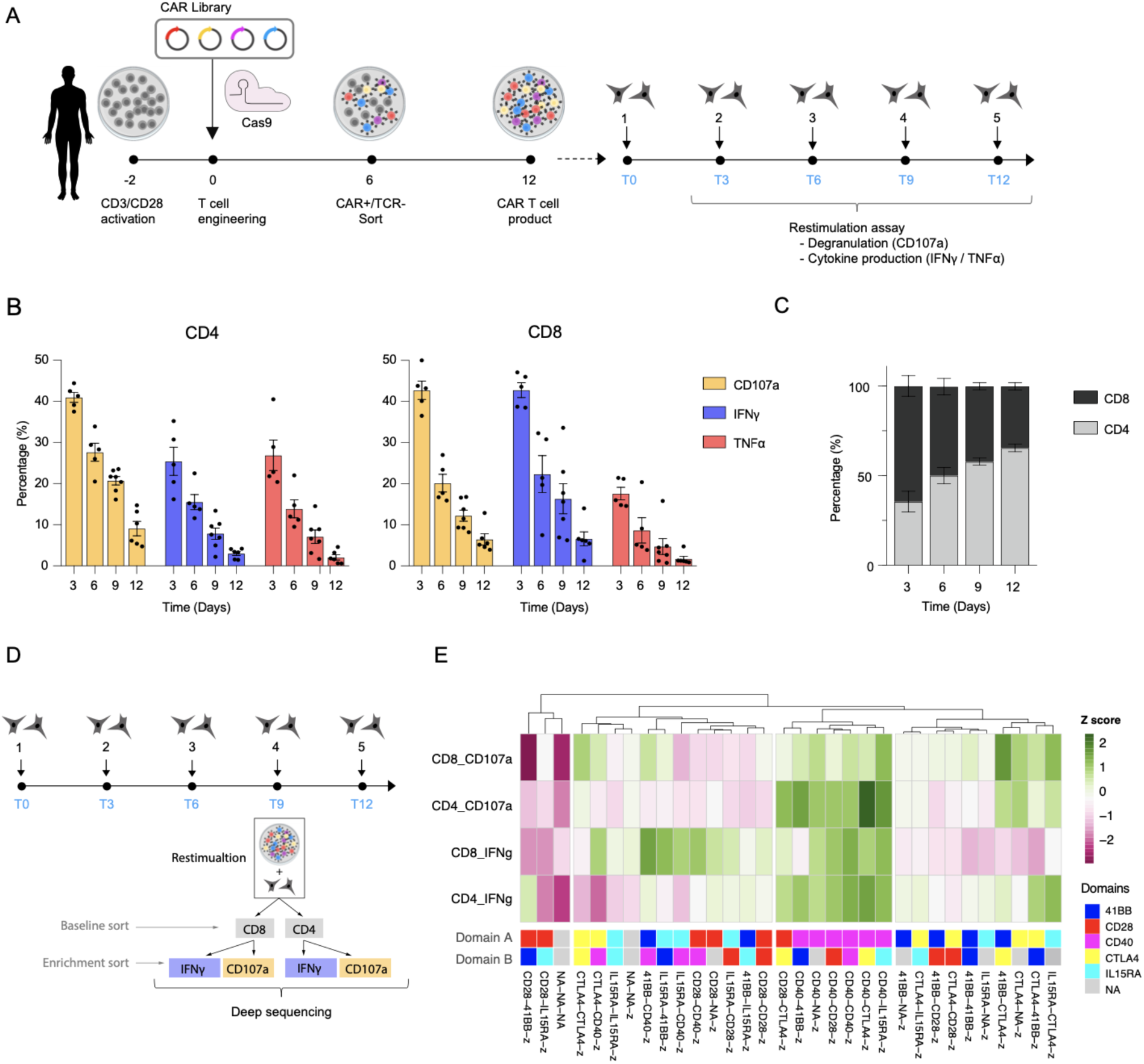
Assessment of T cell dysfunction using a repeated tumor rechallenge assay. **A**- Schematic representing the CAR library T cell production protocol followed by a repeated antigen stimulation (RAS) assay. 12 days after T cell engineering, a purified population of library CAR T cells was co-cultured with HER2+ SKBR3 target tumor cells at a 1:1 effector to target (E:T) ratio. Every three days, the co-culture was reset and a fraction of the cultured cells was used to assess the T cell anti-tumor potential through the RAS assay. **B**- Percentage of CD4 and CD8 cells presenting surface expression of CD107a or intracellular production of IFNγ and TNFα at different stages of the RAS assay. **C**- Percentage of CD4 and CD8 cells observed at different stages of the RAS assay. (**B** and **C**: n=7, technical replicates including 4 different donors. Error bars represent SEM) **D**- Schematic describing the sorting strategy used to assess the enrichment of the CAR library in degranulating (CD107a) and proinflammatory (IFNγ) T cells at a pre-exhausted stage of the RAS assay. **E**- Hierarchical clustering of the CAR library according to the enrichment or depletion of variants following a CD107a or IFNγ positive selection after 9 days of RAS assay. Variants are clustered according to Z-scores, which are calculated based on the log2 fold change in the relative library frequencies before and after enrichment for effector markers shown in panel (D). CD8 and CD4 T cell compartments were analyzed separately (n=3, independent biological replicates). Panels A and D were partially created with BioRender.

Based on the RAS functional characterization, we observed that the library of CAR T cells reached a pre-dysfunctional state by day 9. The anti-tumor potential at this stage was evidently reduced; however, T cells were still able to control tumor cell growth. To assess the persistence of the different CARs, we aimed to resolve the library diversity following a FACS-based selection of cells that remained positive for effector markers (CD107a or IFNγ) by day 9 of the RAS assay (Fig. 2d). Targeted deep sequencing of the CAR transgenes was performed and enrichment scores were computed using post-enrichment library frequencies normalized to baseline (library frequencies on day 9 before selection) for the CD8 and CD4 T cell populations. As expected, the NS-CAR was consistently depleted for every marker (Fig. 2e). Notably, the CD40 signaling domain in position A (proximal to the cell membrane) was a key driver of T cell persistence, resulting in high enrichment scores for all groups (Fig. 2e and Supp. Fig 2). However, CD40 in position B (distal from the cell membrane) showed lower enrichment scores, but still promoted a proinflammatory phenotype in CD8 cells. In addition, CTLA4 in position B was enriched in CD107a+ cells and thus appears to drive a more persistent cytotoxic phenotype. CD28 and 4-1BB signaling domains induced a moderate or reduced persistence.

### Single-cell transcriptional profiling resolves CAR-induced phenotypes

We next sought to further resolve the CAR-induced T cell phenotypes of the library across RAS using the multidimensional readout of scRNAseq. CAR T cell library cells were produced from two healthy donors and transcriptomic data was generated at early, middle and late stages of the RAS assay (days 0, 6 and 12). At each of these time points, CAR T cells were stimulated with HER2-expressing SKBR3 cells for 6h and then sequenced (Fig. 3a). In addition to the scRNAseq data, we performed single-cell cellular indexing of transcriptomes and epitopes (scCITEseq), to detect a panel of T cell surface marker proteins. Finally, we also performed scCARseq using an adapted protocol from our previous work (*16*), which enables de-multiplexing of the pooled CAR library by identifying the CAR variant of each cell. (Supp. Fig. 3).

**Figure 3:**
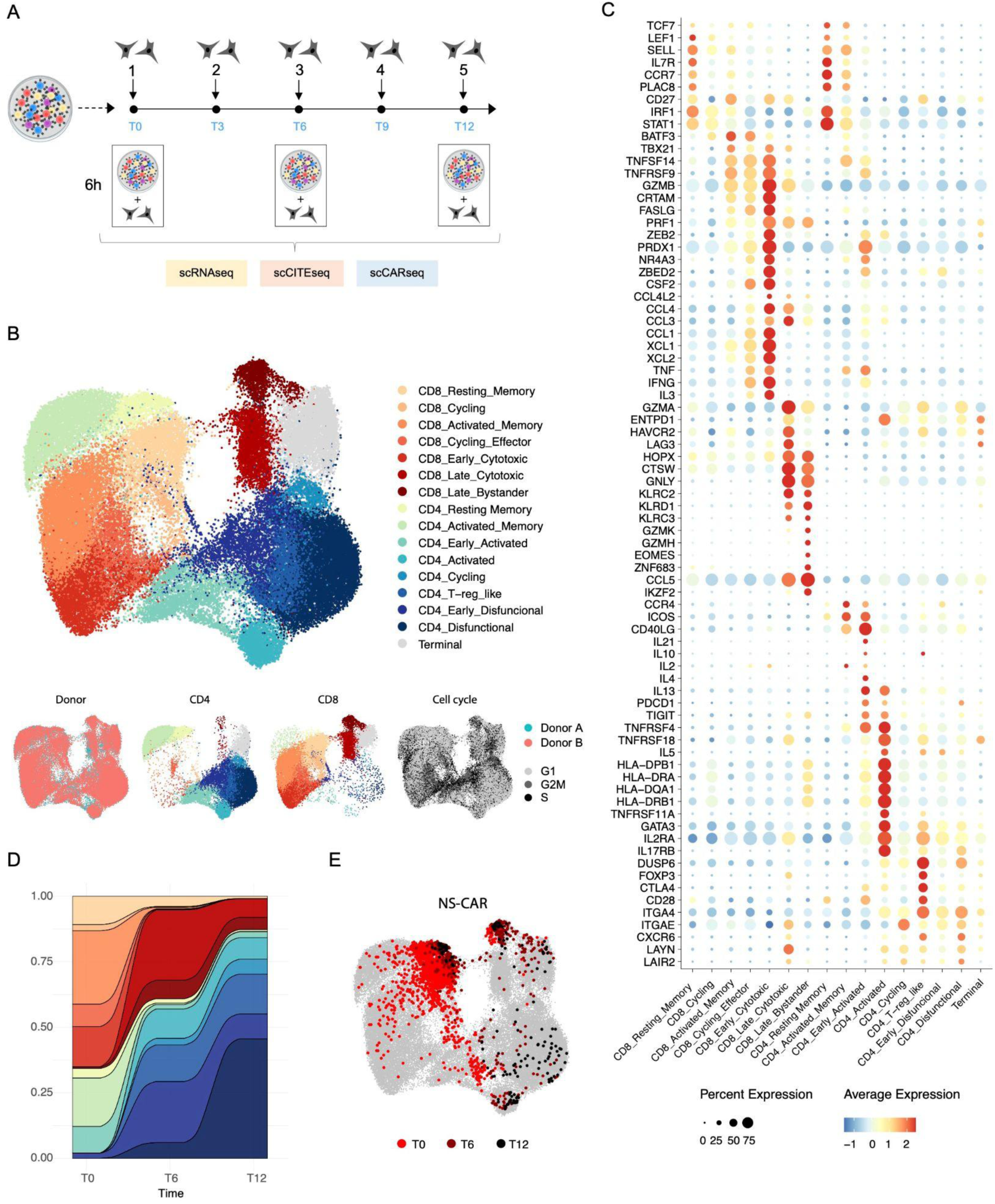
CAR-induced transcriptional phenotypes at single-cell resolution. **A**- Schematic describing the generation of single-cell data. The pooled library of CAR T cells from early, middle and late time points in the repeated antigen stimulation assay (RAS) was stimulated for 6h in the presence of SKBR3 target tumor cells and then processed using scRNAseq, scCARseq and scCITEseq. **B**- UMAP embedding and unsupervised cell clustering of the scRNAseq data generated as described in (**A**). 58,949 cells from 2 healthy donors and three time points are shown. At the bottom, UMAP embeddings are colored based on Donor, CD4 or CD8 annotation and cell cycle phase. **C**- Dot plot shows the expression of a selection of differentially expressed T cell marker genes that are used to annotate the clusters described in (**B**). **D**- Change in cluster representation across time points. **E**- Distribution of cells annotated to display a non-signaling CAR (NS CAR) within the UMAP embedding of (**B**). Panel (**A**) was partially created with BioRender.

Single-cell sequencing and data processing resulted in a total of 62,934 annotated CAR T cells across the three different time points, with full coverage of the CAR library across every time point and donor. An additional random subsampling of abundant variants was performed to correct for arbitrary clonal expansion, resulting in 58,949 cells, which were used for downstream phenotypic analysis. The lack of correlation between the expression of TCR variable genes across CAR variants, time points or donors validated the presence of sufficient clonal diversity in our library (Supp. Fig. 4). Dimensionality reduction by uniform manifold approximation and projection (UMAP) and unsupervised cell clustering separated cells into 16 different clusters (Fig. 3b). Annotation of the clusters was based on CD4 and CD8 expression, cell cycle phase prediction and differential gene and surface expression of key T cell marker genes (Fig. 3c and Supp. Fig. 5); both CD4 and CD8 T cell subsets presented a resting memory cluster characterized by the expression of TCF7, CCR7, LEF1 and SELL genes and protein surface expression of CD45RA and CD62L. The CD8 memory cluster then progressively transitioned into activated and effector phenotypes characterized by the increased expression of activated (TNFRSF9, TBX21, ZBED2), cytotoxic (GZMB, PRF1, FASLG) and proinflammatory genes (CRTAM, IFNG, TNF, CSF2, XCL1, XCL2) and eventually into a late cytotoxic phenotype characterized by the expression of late effector differentiation genes such as HOPX, ENTPD1, LAG3, HAVCR3 and GNLY. Cytotoxicity was also evidenced by the increased surface detection of HER2 on CAR T cells as a result of trogocytosis, a process by which there is a unidirectional transfer of plasma membrane and associated surface proteins from target cells to effector lymphocytes (*26*) (Supp. Fig. 5c). Lastly, a CD8 cluster was observed that presented a CAR-independent, bystander T cell signature (GZMK, GZMH and several KLR genes), which could be attributed to the effect of the cytokine storms and the cell killing environment on unstimulated T cells. Likewise, the CD4 memory cluster also transitioned into activated and more differentiated phenotypes evidenced by the expression of activation genes such as CD40LG, IL2RA, ICOS, TNFSF14, TNFSF and IL17RB and a broad range of cytokines. This activated phenotype later transitioned into a rather dysfunctional phenotype and a Treg-like cluster characterized by the expression of FOXP3 and CTLA4. Lastly, a mixed CD4 and CD8 cluster, high in mitochondrial gene expression, was annotated as a terminal phenotype.

The progression of T cell phenotypes from a memory and early activation state, through a potent effector phenotype, to a late, less functional state correlated with the scCITEseq data for surface expression of early, middle and late T cell activation markers (Supp. Fig. 5b), as well as the time points at which the samples were collected (Fig. 3d and Supp. Fig. 6a). As previously observed, the CD8 compartment was drastically reduced through RAS progression in favor of a growing ratio of dysfunctional CD4 CAR T cells. The absence of CD8 cells presenting a terminally exhausted phenotype and the drop in the overall number of T cells in late co-cultures suggest the death of CD8 cells following their terminal effector differentiation.

Having resolved the recorded T cell phenotypes, scCARseq enabled us to de-multiplex the CAR library identity and investigate how different CAR signaling architectures can drive distinct T cell responses upon both initial and repeated antigen stimulation. First, we examined the T cell phenotypes of the NS-CAR through time (Fig. 3e). As expected, the lack of CAR signaling domains resulted in non-activated T cells that remained in a resting memory phenotype at early and even late time points, with only a small fraction of cells presenting a bystander T cell activation phenotype. Cluster enrichment of CD4 and CD8 cells across time points for the rest of the library variants indicated that every other CAR was able to trigger T cell activation, as evidenced by the lack of cells presenting a resting memory phenotype (Supp. Fig. 6b).

### Role of signaling domain combinations in early activation of CAR T cells

To understand how signaling domain combinations shape the early activation of T cells, we examined transcriptional phenotypes after 6h of tumor co-culture. For both CD8 and CD4 subsets, we could observe the separation of cells across a T cell differentiation axis. When ordering cells according to a predicted pseudotime, the annotated clusters indeed followed such a trajectory, evolving from a resting memory to a potent effector phenotype (Supp. Fig. 7). The enrichment of CAR variants across these clusters can therefore reveal differences in early activation signatures triggered by the different CARs. For the CD8 cell compartment, the presence of the CD40 domain in position A appeared to be the main driver of a fast and potent effector phenotype, as all CD40 variants (except CD40–41BB) presented the highest percentage of cells within the effector and cytotoxic clusters (Fig. 4a). On the other hand, 4-1BB containing CARs, while still activated, appeared to trigger a less potent but stronger effector memory-like phenotype. Notably, CD4 cells showed a different trend; for example, CTLA4-containing CARs appeared to drive the strongest CD4 activation and differentiation, while CD40, CD28 and 4-1BB retained an overall CD4 effector memory phenotype.

**Figure 4:**
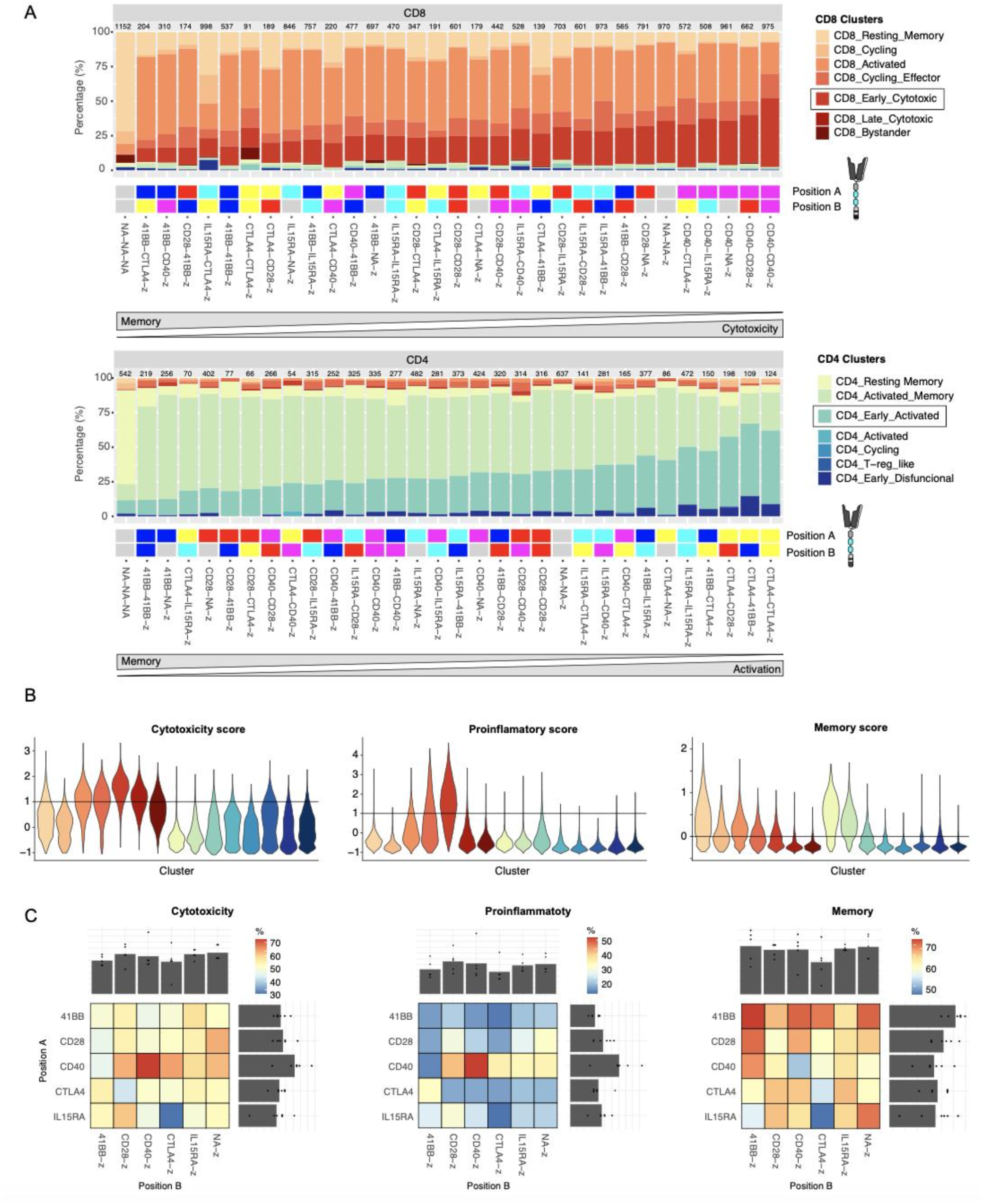
CAR-specific signatures following initial CAR T cell activation. **A**- Cluster enrichment observed for the different CAR T cell variants following 6h of tumor co-culture. CD8 and CD4 annotated cells are shown in different plots and variants are ordered (right to left) by the enrichment in the cluster highlighted with a black box. Under each bar plot, a heatmap describes the intracellular domain combination of the library candidates. The number of cells used to define the cluster distribution is reported at the top of each bar. **B**- Distribution of single-cell gene-set scores across the different clusters described in Figure 3. A horizontal line determines a threshold at which cells are considered to be positive for each given score. **C**- Heatmaps show the percentage of cells with a positive score based on the thresholds described in (**B**) following 6h of tumor co- culture. Each heatmap separates variants based on the CAR signaling domains in position A (proximal to the cell membrane) or position B (distal from cell membrane). In addition, bar plots at the top and right-hand side of the heatmap compile the frequencies for all variants presenting a given domain in the different positions. Heatmaps for cytotoxicity and proinflammatory scores only include CD8 cells, while the memory score includes both CD4 and CD8 cells.

In addition to cluster enrichment, we used single-cell gene set scoring to further resolve the activation signatures based on the simultaneous expression of several marker genes. The CD8 effector phenotype was assessed for its cytotoxic and proinflammatory potential, and a memory phenotype score was computed for all cells (Fig. 4b). Based on these scores, in silico sorting of cells was performed to assess the different CAR library variants by their enrichment in such phenotypes. Using the NS-CAR to set a baseline threshold, we then investigated the impact of CAR signaling domain composition on the appearance of each of these phenotypes (Fig. 4c). As described previously, all CAR constructs were able to trigger a strong cytotoxic phenotype (40-70% of CD8 cells); however, once again, the CD40 domain in position A appeared to drive a particularly high cytotoxicity that was enhanced when the CD40 domain is repeated. This pattern is even more striking when examining the proinflammatory signature. CD40 in position A also resulted in the most powerful proinflammatory phenotype that appears to be slightly restrained when incorporating 4-1BB or CTLA-4. The second generation 28z CAR, as expected, induced several of the most potent cytotoxic and proinflammatory signatures, serving as validation of our results. Lastly, the memory phenotype signature seemed to be highly enriched in 41BB-containing CARs, once again aligning with previous studies (*27*, *28*).

### CAR co-stimulation can modulate long-term T cell persistence

A common limitation often faced by CAR T cell therapies is the transient persistence of T cells, ultimately resulting in their inability to control tumor growth and disease progression (*4*, *29*). Identifying CAR design features that promote a more persistent phenotype is of substantial value. To address this, we next leveraged the RAS assay to study the progression of T cell phenotypes across the CAR library. The scRNAseq data of CAR T cells from middle and late time points in the RAS assay were separated by CD4 and CD8 annotation and re-clustered to further resolve the RAS late-stage phenotypes.

Amongst the CD8 compartment, we observed two clusters, annotated as proinflammatory (CRTAM, IFNG, CSF2) and cytotoxic (PRF1, GZMB, GNLY, IL2RA) that still present effector potential (Fig. 5 a-c). Excluding a resting memory-associated cluster, the remainder of the clusters start to lose the expression of key effector marker genes, reaching a dysfunctional and subsequent terminal phenotype that ultimately leads to cell death. Using the enrichment of proinflammatory and cytotoxic clusters at a RAS late time point (12 days) as a marker for persistence, we observed that CD40, mainly in position A, appeared to be a key domain for long-term persistence (for both proinflammatory and cytotoxicity phenotypes). 4-1BB and CTLA4 promoted a late-stage cytotoxic, but not a proinflammatory, phenotype. IL15RA and CD28, on the other hand, had the largest percentage of cells already transitioning into a dysfunctional phenotype (Supp. Fig. 8).

**Figure 5:**
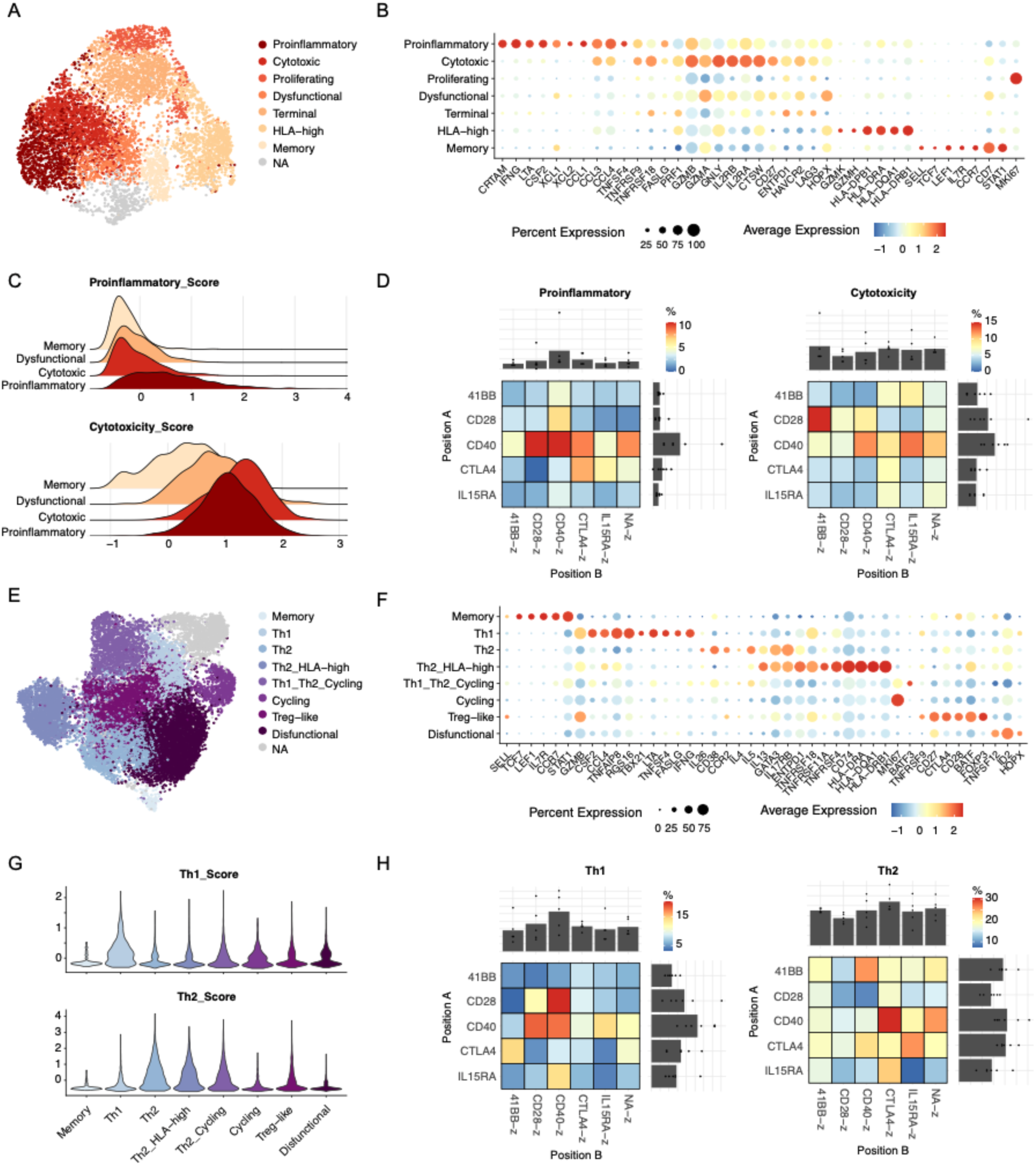
CAR-specific signatures following repeated antigen stimulation. **A**- UMAP embedding and unsupervised cell clustering of CD8 annotated cells at middle and late time points in the RAS assay (6 and 12 days). Cluster annotated as NA includes misannotated cells presenting a CD4-specific phenotype. **B**- Dot plot shows the expression of a selection of differentially expressed T cell marker genes, used to annotate the clusters described in (**A**). **C**- Distribution of single-cell gene-set scores across relevant clusters described in (**A**). **D**- Heatmaps show the enrichment of CD8 cells in the proinflammatory or cytotoxic clusters at a late time point in the RAS assay (12 days). The enrichment is corrected by the CD8/CD4 ratio fold change from day 0. Each heatmap separates variants based on the CAR signaling domains in position A (proximal to the cell membrane) or position B (distal from the cell membrane). In addition, bar plots at the top and right-hand side of the heatmap compile the frequencies for all variants presenting a given domain in the different positions. **E**- UMAP embedding and unsupervised cell clustering of CD4 annotated cells at middle and late time points in the RAS assay (6 and 12 days). Cluster annotated as NA includes misannotated cells presenting a CD8-specific phenotype and dying cells with high mitochondrial gene expression. **F**- Dot plot shows the expression of a selection of differentially expressed T cell marker genes that are used to annotate the clusters described in (**E**). **G**- Distribution of single-cell gene-set scores across different clusters described in (**E**). **H**- Heatmaps show the enrichment of CD4 cells in the Th1 or Th2 clusters at a late time point in the RAS assay (12 days). Each heatmap separates variants based on the presence of CAR signaling domains in position A or position B . In addition, bar plots at the top and right-hand side of the heatmap compile the frequencies for all variants presenting a given domain in the different positions.

Another feature to take into consideration within the CD8 T cell compartment is the decline in CD8 cell numbers over time. As previously mentioned, this reduction in the CD8/CD4 ratio appears to correlate with terminal effector differentiation, ultimately leading to cell death of only CD8 cells. A faster drop in CD8/CD4 ratio can therefore be associated with a lack of persistence. By combining CD8/CD4 fold change (Supp. Fig. 9) with the enrichment in the late effector clusters, we can obtain a more comprehensive persistence score (Fig. 5d). Based on this, CD40 once again proves to be the signaling domain that induces the most persistent phenotype, followed by CTLA-4.

Amongst the CD4 subset, another two main functional clusters, a proinflammatory Th1 (TBX21, CRTAM, IFNG, GZMB) and a polyfunctional Th2 cluster (GATA3, IL4, IL5, IL13), were identified in addition to other cycling, T- reg-like and dysfunctional clusters (Fig. 5e and f). The Th1 and Th2 signature was also confirmed by gene set scoring (Fig. 5g). Cluster enrichment was then used to evaluate the persistence of CD4 CAR T cell variants. CD40 consistently

drove the most persistent proinflammatory signature by being the most enriched in the Th1 cluster, while CTLA-4 and 4-1BB promoted a Th2-enriched persistent phenotype (Fig. 5h). On the other hand, CD28-containing CARs were consistently the most enriched variants in the dysfunctional cluster, suggesting once again that CD28-containing CARs are prone to induce poor persistence (Supp. Fig. 8).

## DISCUSSION

Domain recombination has been pivotal in the evolution of signaling networks, operating on the principle that the function of a protein domain is modular and can promote new functions when embedded differently within a cellular network (*30*). As a product of rational domain recombination, CARs can be further optimized through additional signaling domain rearrangements. Despite the remarkable progress in the field of CAR T cell engineering (*5*, *31*), significant gaps persist in understanding how changes in the CAR signaling architecture impact resulting T cell phenotypes and their therapeutic potential. In particular, costimulatory domains have proven to be key in providing CAR T cells with essential properties for clinical efficacy, but the impact of changing the type, number and order of costimulatory domains has yet to be systematically characterized. In this study, we bridge these gaps by combining the use of a combinatorial CAR signaling domain library, pooled screening assays and scRNAseq to systematically study the dynamics governing CAR-induced phenotypes at high resolution.

We evaluated the impact of recombining five immune-receptor signaling domains, resulting in a 32-candidate CAR library that revealed architecture-specific patterns. While all CAR designs were capable of eliciting a T cell activation response, the incorporation of CD28 or 4-1BB domains replicated known phenotypic features associated with these domains (*27*, *28*, *32*); CD28 induced a potent but less persistent T cell activation, while 4-1BB promoted an effector memory phenotype. In parallel comparisons with benchmark CARs, insights into three additional domains were observed. CD40 consistently distinguished itself by triggering the most potent and persistent T cell responses. This aligns with previous findings indicating that CARs combining the signaling domains of MyD88 or CD79A with CD40 exhibit superior proliferation and antitumor activities in preclinical tumor xenograft mouse models (*6*, *20*). Moreover, CD40-containing CARs were selected amongst the top candidates when performing pooled screens of two CAR libraries (*13*, *14*), further highlighting the role of this domain in enhancing CAR signaling.

CTLA4, recognized as an inhibitory receptor on T cells, when embedded in the CAR signaling architecture resulted in potent CARs capable of promoting a robust T cell response, particularly amongst the CD4 compartment, with a persistent cytotoxic phenotype. Despite the inhibitory effect of CTLA4 signaling (*23*), our findings align with recent research reporting that the addition of CTLA4 cytoplasmic tails to a 28z CAR led to increased cytotoxicity and reduced production of pro-inflammatory cytokines (*7*). Despite its previous association with enhancing T cell anti-tumor potential (*21*), the IL15RA cytoplasmic domain did not seem to provide impactful T cell co-stimulation amongst the variants. The reduced size of its intracellular domain, coupled with the central role of its extracellular portion in carrying out its molecular function (*33*), suggests that the IL15RA domain may act as a molecular spacer within the CAR architecture.

With regard to domain positioning, our study also revealed distinctive patterns. In agreement with prior research that addressed the impact of altering the positioning of domains within the CAR (*34–36*), we observed that domains located closer to the cell membrane exhibited a dominant effect on phenotype. For instance, CD40 and 4-1BB respectively induced a distinct polarization towards effector or memory phenotype mainly when present in a membrane-proximal position. Nevertheless, domains in a more distal position continued to exert influence on phenotype, seemingly producing an additive effect. For example, CD40-z demonstrated a potent cytotoxic and proinflammatory phenotype, further potentiated in CD40-CD40-z but moderated in CD40-4-1BB-z, favoring a more memory-like phenotype. Despite these general observations, the mechanistic complexity associated with the introduction of domain rearrangements within a signaling network is far more sophisticated and highly dependent on the nature of the domains used. For example, CTLA4 displayed a more prominent role in promoting cytotoxicity when situated in a membrane-distal position. This observation, also suggested by Zhou et al. (*7*), may be linked to the role of its endocytosis motif in receptor recycling. This mechanistic feature could benefit from distal positioning, while distancing CTLA4 inhibitory signaling from the dominating membrane-proximal position.

The enormous complexity associated with rewiring signaling networks highlights the value of conducting systematic studies of CAR signaling domain rearrangements. In addition, the diversity of the CAR signaling domain combination space requires high-throughput approaches that enable parallel comparisons of multiple architectures. Pooled screening of CAR libraries combined with scRNAseq provides such a high-throughput approach (*13*, *14*, *16*, *37*). An exciting frontier of this field is the integration of CAR libraries with machine learning, as previously demonstrated by Daniels et al. (*15*). Machine learning-guided CAR T cell engineering may further elucidate mechanistic nuances of signaling domains and enable novel CAR designs. The compact yet systematic design of our library, combined with the comprehensive and high-resolution data generated in this study, may provide training data for machine learning models that are able to decipher the rules of CAR signaling.

While our study provides valuable insights into the intricate landscape of CAR signaling and its impact on T cell phenotypes, we acknowledge certain limitations and outline future perspectives. In the context of pooled screens, the unavoidable bystander effect resulting from paracrine signaling poses logical concerns. Despite this limitation, the inclusion of a NS CAR as a negative control validated the minimal impact of this paracrine effect in overall T cell phenotype and stresses the importance of incorporating such controls in pooled library assays. Secondly, reproducing a clinically relevant T cell activation context poses a significant challenge. In vitro co-culture lacks the cellular heterogeneity, tumor microenvironment and anatomical barriers encountered in real clinical scenarios. While in vivo settings attempt to address some of these challenges, human tumor xenograft mouse models in immunocompromised mice often also fall short in replicating clinical conditions. Our choice of using ex vivo RAS provided extensive phenotypic characterization of CAR signaling in a simplified setup mimicking the clinical challenge of CAR T cell dysfunction following chronic antigen exposure. Despite this, the limited understanding of the correlation of CAR T cell phenotypes with clinical outcomes still makes it difficult to speculate which variant could exhibit better clinical performance, necessitating further functional validation (*38*). Lastly, while this study provides a systematic and in-depth analysis of such a large number of CARs, such an approach can be adapted to larger libraries or can include other CAR modules, such as antigen binding domains of varying affinities (*39*), hinges and transmembrane domains (*40*).

## MATERIALS AND METHODS

### Library cloning

The CAR library was cloned using a Type II restriction enzyme cloning strategy as previously described (*41*). A backbone plasmid containing an anti-HER2 first generation CAR gene (composed of a CD8ɑ secretion peptide, a Herceptin-derived scFv (4D5), two Strep tags, CD28 hinge and transmembrane domains, the CD3ζ cytoplasmic region and a bGH polyA sequence) flanked by *TRAC* locus-specific homology arms was generated. In addition, a cloning cassette with inverted AarI sites was introduced between the TMD and the CD3ζ sequence. Lastly, a 3’ UTR barcode sequence was added using an overhang PCR and recircularization strategy. Synthetic gene fragments containing the cytoplasmic sequence of CD28, 4-1BB, CD40, IL15RA and CTLA4 genes were generated (Twist Bioscience) with different sets of flanking sequences containing an AarI recognition site that allows for the ligation of a defined number and order of domains within the CAR backbone. Domain sequences were individually amplified, digested with AarI (Thermo Fisher; 4 h, 37 °C) and ligated into the previously digested backbone plasmid using a T4 ligase (NEB; 30 min, 37 °C). For each library candidate, the ligated plasmids were transformed into E. coli DH5ɑ cells, purified and sequence verified using Sanger sequencing. The NS-CAR and the 1st generation CAR were independently cloned using deletion Q5 mutagenesis. Finally, all library candidate plasmids were pooled at a 1:1 ratio.

### Primary human T cell isolation and culture

Buffy coats from healthy donors were acquired through the Blutspendezentrum Basel (University of Basel). All participating volunteers provided written informed consent in accordance with the general guidelines approved by Swissethics (Swiss Association of Research Ethics Committees). Peripheral blood mononuclear cells (PBMCs) were isolated using a Ficoll-based density gradient and stored in liquid nitrogen until needed. Immediately after thawing, negative selection of T cells was performed using the EasySep human T cell isolation kit (Stemcell) and cultured in X-VIVO 15 medium (Lonza) supplemented with 5% fetal bovine serum (FBS), 50 μM β-mercaptoethanol, 50 μg/mL Normocin (Invivogen) and 100 U/mL IL-2 (Peprotech), referred to as T cell growth medium.

### Primary human T cell genome editing

Primary human T cells were engineered to integrate a CAR gene into the *TRAC* genomic locus using CRISPR-Cas9 genome editing. Double-stranded DNA HDR template and Cas9 ribonucleoprotein (RNP) were prepared as previously described (*16*). T cells were activated using Human T-Activator CD3/CD28 Dynabeads (Thermo Fisher) at a 1:1 cell:bead ratio in T cell growth medium. After 48 h, beads were magnetically removed and cells were electroporated using the Lonza 4D electroporation system. To do this, 1x10^6^ cells were washed once in PBS, resuspended in 20 uL of P3 nucleofection buffer (Lonza) containing 1.2 uL of Cas9 RNP mix and 0.4 ug double-stranded DNA HDR template and electroporated using the EH-115 program. After electroporation, cells were immediately recovered in 150 uL of T cell growth medium. For each batch of CAR library T cells, at least 1x10^7^ T cells were engineered to achieve sufficient clonal diversity across all candidates.

### Cell line culture

SKBR3-GFP cell line was cultured in Dulbecco’s Modified Eagle Medium (DMEM; Gibco) supplemented with 10% FBS, 1% penicillin-streptomycin (Gibco) and 50 μg/mL Normocin (Invivogen).

### CAR T cell staining and cell sorting

Flow cytometry was used to analyze and select correctly engineered CAR T cells based on the positive staining of a StrepTag located in the extracellular portion of the CAR and the lack of expression of the TCR complex. A two-step staining strategy was employed, initially using a biotinylated anti-Strep tag antibody (Supp. Table 3), followed by a combination of streptavidin-BV421 conjugate and CD3ε-APC antibody (Supp. Table 3). T cells were washed in FACS buffer (PBS, 2 % FBS, 1 mM EDTA) and then incubated in FACS buffer containing the antibody mix for 20 minutes. Cells were then washed again and analyzed using a Fortessa LSR flow cytometer (BD) or sorted using a FACS Aria Fusion (BD).

### Deep sequencing of CAR libraries

The diversity of library CAR T cells was determined using deep sequencing. Genomic DNA from 5,000 - 50,000 CAR-expressing T cells was extracted using Quick Extract (Lucigen) and used as the template for a 2-step PCR strategy. In a first PCR reaction, primers F1 and R1 (Supp. Table 2) were used to amplify a region of the CAR gene (2000 - 2500 bp), which confirmed the CAR integration into the *TRAC* locus. Following a 0.6X SPRIselect bead DNA cleanup (Beckman Coulter) the DNA product was used as a template for a second PCR reaction using primer mix F2 and R2 (Supp. Table 2). This amplified a 261 bp sequence in the CAR 3’UTR region that contained the barcode sequence, which determines its library identity. The resulting amplicons were purified using a 1.2X-0.6X double-sided SPRIselect bead DNA cleanup (Beckman Coulter), prepared for sequencing using a KAPA HyperPrep kit (Roche) and sequenced on an Illumina MiSeq system. Sequencing data analysis was performed using the Biostrings package in R.

### In vitro repeated antigen stimulation (RAS)

To simulate a chronic antigen stimulation, CAR-T cells were repeatedly co-cultured with the HER2-expressing tumor cell line SKBR3. On day 14 (12 days after bead removal and T cell engineering) T cells were co-cultured with SKBR3- GFP cells at a 1:1 E:T ratio in CAR media supplemented with 30 IU/mL of IL-2. Every 3 days cells were counted using a hemocytometer and new SKBR3-GFP cells were added to re-adjust the co-culture to a 1:1 E:T ratio.

### Degranulation and cytokine production assay

To assess the effector potential of co-cultured CAR-T cells, we measured degranulation and the production of cytokines following the restimulation of CAR-T cells with SKBR3-GFP cells. 50,000 CAR-T cells were co-cultured with 100,000 target cells for 5h in the presence of CD107a antibody (Supp. Table 3) and 1x Brefeldin A (Biolegend). Following co-culture, cells were stained for dead cells (Zombie NIR; Biolegend) and surface markers (CD4 and CD8; Supp. Table 3), fixed using Fixation Buffer (Biolegend) and stained for the intracellular accumulation of IFNγ and TNFα (Supp. Table 3) in 1x Permeabilization Buffer (Biolegend). Samples were analyzed using a Fortessa LSR flow cytometer (BD) or sorted using a FACS Aria Fusion (BD).

### Single-cell sequencing (scRNAseq)

Library CAR T cells derived from two healthy donors were subjected to a RAS assay, as previously described. At days 0, 6 and 12 of the RAS assay, CAR library T cells were co-cultured with SKBR3-GFP cells for 6h in 30 IU/mL of IL-2. Following this time, the co-cultures were washed with FACS buffer, stained using DRAQ7, and the live GFP-negative population was sorted using a FACS Aria Fusion (BD). Cells were then stained using 20 Totalseq B antibodies (Supp. Table 4) and introduced into the Chromium Single Cell 3’ scRNAseq pipeline v3.1 (10x genomics) following the manufacturer’s guidelines (User guide CG000317 Rev D). In short, 20,000 cells were loaded into each Chromium chip lane to generate single-cell emulsions containing barcoded oligonucleotides that allow the generation of barcoded cDNA from mRNA and oligo-tagged antibodies. Using the amplified cDNA as a template, scRNAseq and scCITEseq libraries were generated and sequenced using the Illumina Novaseq platform.

### Single-cell CAR sequencing (scCARseq)

Demultiplexing of the CAR library to define the CAR identity for each cell in scRNAseq data was achieved using an adapted version of a previously described scCARseq methodology (*16*) (Supp. Fig 3). Using 10 uL of the cDNA product resulting from the single-cell sequencing pipeline, the 3’ UTR region of the CAR transcripts, containing a CAR variant specific barcode (CAR-BC), was amplified using F3 and R3 primers (Supp. Table 2) and KAPA-Hifi polymerase (Roche). Following a 1X SPRIselect bead DNA cleanup (Beckman Coulter), the DNA product containing partial Illumina-specific adaptors was further amplified and indexed using Dual Index Kit TT, Set A primers (10x Genomics, PN-1000215). The final scCARseq library was then purified using a 1X-0.6X double-sided SPRIselect bead DNA cleanup (Beckman Coulter) and sequenced with the Illumina platform using the same cycle scheme as the scRNAseq and scCITEseq libraries. scCARseq data analysis was conducted using the Biostrings package in R. Only cells with at least 2 different unique molecular identifiers (UMIs) defining the same CAR annotation were accepted.

### Single-cell sequencing data analysis

The raw sequencing data was aligned to the GRCh38 human reference genome using Cell Ranger (10x Genomics, version 6.0.0) and imported into R (version 4.2.3) to perform downstream analysis using the Seurat package (version 4.3.0.1). In the first place, only cells assigned to a single CAR variant were selected. Low-quality cells were removed based on the detection of a low number of genes (*nFeature_RNA > 300*), high number of gene expression UMIs (*nFeature_RNA < 50,000*), high number of Antibody-Derived Tags (ADT) UMIs (*nCount_ADT < 30,000*), or a high percentage of mitochondrial genes (*percent.mt < 20*). Lastly, in order to correct for arbitrary clonal expansion that may occur through RAS, a subsampling step was performed; for each sample (different time point or donor), CAR variants exceeding two times the theoretical balanced library distribution (maximum 6.25% of cells per CAR variant) were randomly subsampled to meet this criteria.

The resulting 58,949 single-cell transcriptomes were then normalized, scaled while regressing out the effect of cell cycle phase and percent of mitochondrial genes and finally integrated using Harmony (*42*) (applying a lambda of 1 and 200 for sample variables ‘Donor’ and ‘Time’ respectively). Dimensionality reduction using UMAP and unsupervised cell clustering was then used to visualize and analyze the resulting T cell phenotypes. ADT data was normalized using dsb (*43*) in *Python*, using the parameters ‘pseudocount=10’ and ‘denoise counts=True’. Empty droplets were estimated by dsb from the raw output of Cell Ranger after the exclusion of the cell-containing barcodes found in the filtered output. RNA and ADT data were combined in the annotation of cells as CD4 or CD8. The Seurat object was then further split by CD4/CD8 subsets and time point to perform a more resolved transcriptomic analysis. This analysis included the use of a single-cell gene set scoring function from the Seurat package (AddModuleScore) using the gene sets in Supp. Table 5 and pseudotime and trajectory analysis using the Monocle3 package (*44*).

## ACKNOWLEDGMENTS

We thank Dr. Raphaël B. Di Roberto, Florian Bieberich, Fabrice S. Schlatter, Anna Mei and Kevin Letscher for valuable scientific discussions. We thank the ETH Zurich D-BSSE Single Cell facility and the ETH Zurich D-BSSE Genomics facility for their excellent support and assistance throughout this study. This work was supported by NCCR Molecular Systems Engineering, Switzerland (to S.T.R.); Personalized Health and Related Technologies (to S.T.R.) and the IBM Exploratory Challenge (*AI-driven engineering of the immune system*). A.D. received funding from the European Union’s Horizon 2020 research and innovation programme under the Marie Skłodowska-Curie grant agreement No 955321. J.A.F. received funding from the ThinkSwiss Research Scholarship.

## AUTHOR CONTRIBUTIONS

R.C.R. and S.T.R. designed the study; R.C.R., K.K.W., J.L.F. and J.A.F. performed experiments; R.C.R., K.K.W. and A.D. performed and interpreted bioinformatic analyses; M.R.M provided bioinformatics expertise; R.C.R. and S.T.R. discussed results and R.C.R. and S.T.R. wrote the manuscript with input and commentaries from all authors.

## DATA AND MATERIALS AVAILABILITY

All data are available in the main text or the supplementary materials. Additional data files and code that supports the findings of this study is available from the corresponding authors upon reasonable request. The raw and processed single-cell sequencing data generated in this study will be deposited in the Gene Expression Omnibus.

