## Supplementary Material for "Dissecting the role of CAR signaling architectures on T cell activation and persistence using pooled screening and single-cell sequencing"

SUPPLEMENTARY INFORMATION:

SUPPLEMENTARY FIGURES 1-10

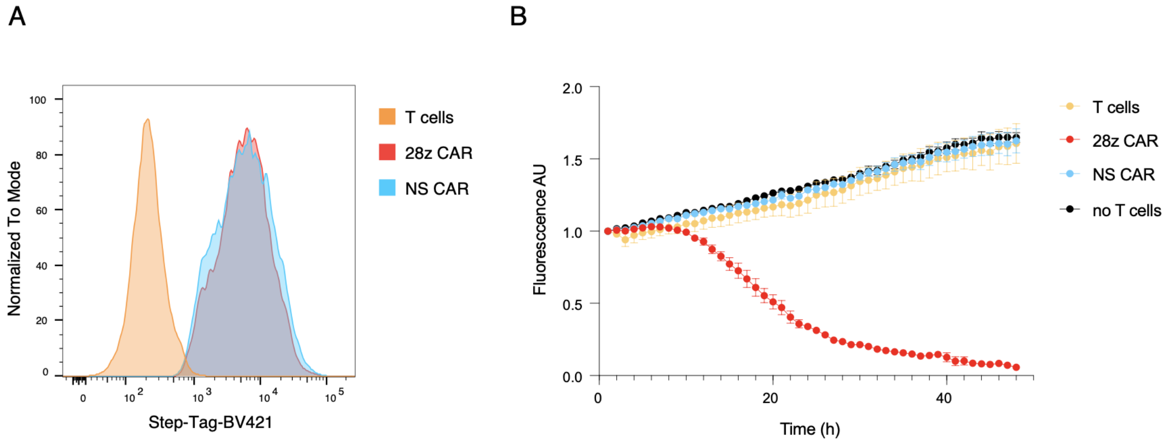

**Supp. Figure 1: A non-signaling CAR serves as an internal library negative control**

**A**- Overlaid histograms show the CAR surface expression profiles of 28z and NS CAR T cells compared to unedited T cells. An antiStrep-Tag-BV421 antibody was used to detect CARs by flow cytometry.

**B**- CAR T cell mediated cytotoxicity of 28z and NS CAR T cells compared to unedited T cells. T cells and HER2+GFP+ SKBR3 cells were co-cultured at a 1:1 E:T ratio and the change in GFP fluorescence was monitored over time (n=3, technical replicates, error bars represent SEM).

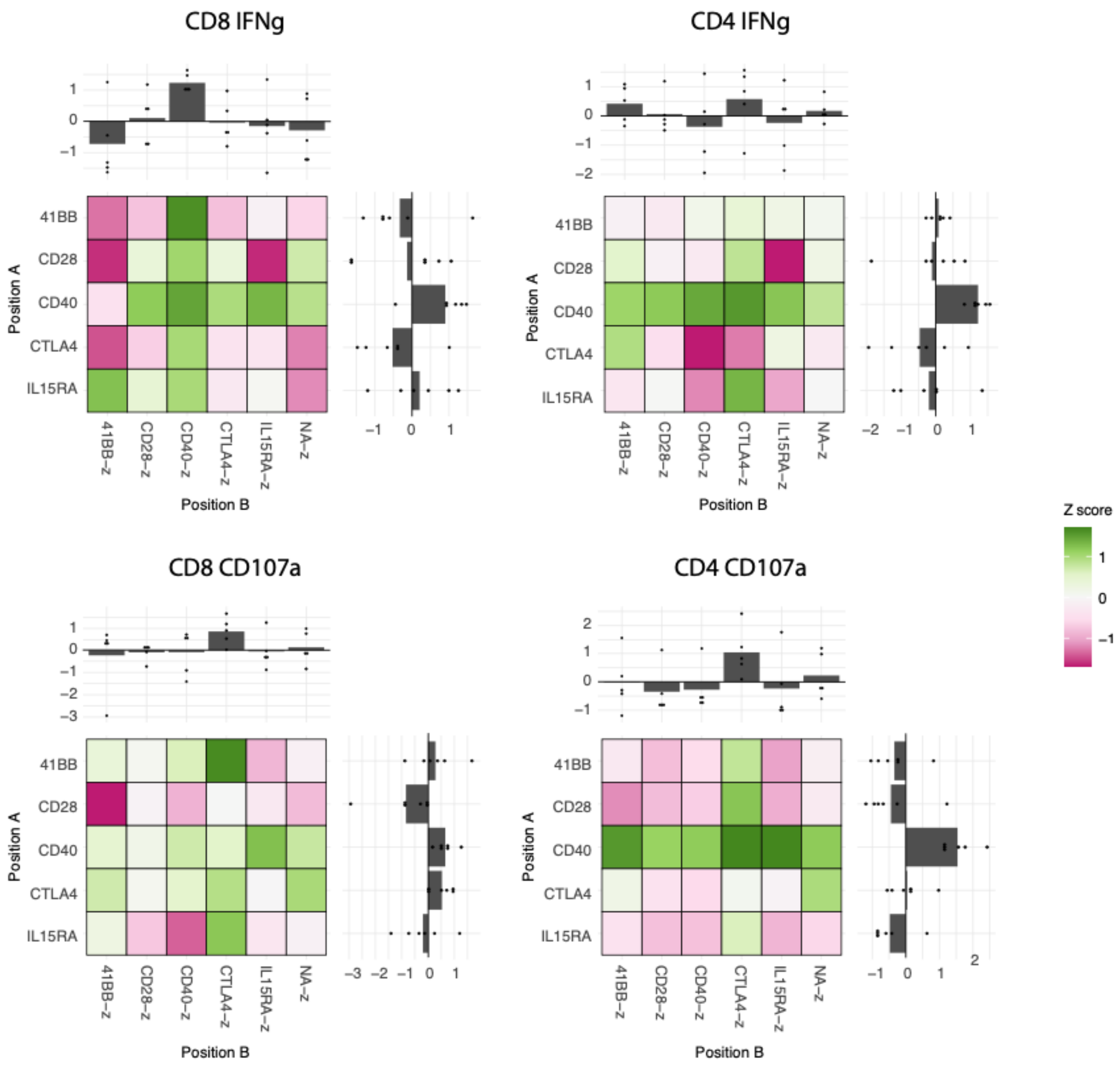

**Supp. Figure 2: Domain identity and position impact CAR T cell persistence during repeated antigen stimulation**

Heatmaps show the enrichment or depletion of CAR variants following a CD107a or IFNγ positive selection after 9 days of RAS. Z-scores were calculated based on the fold change in relative library frequencies before and after selection. CD8 and CD4 T cell compartments were analyzed separately. Each heatmap separates variants based on the presence of CAR signaling domains in position A (proximal to the cell membrane) or position B (distal from cell membrane). In addition, bar plots at the top and right-hand side of the heatmap compile the scores for all variants presenting a given domain in a given position.

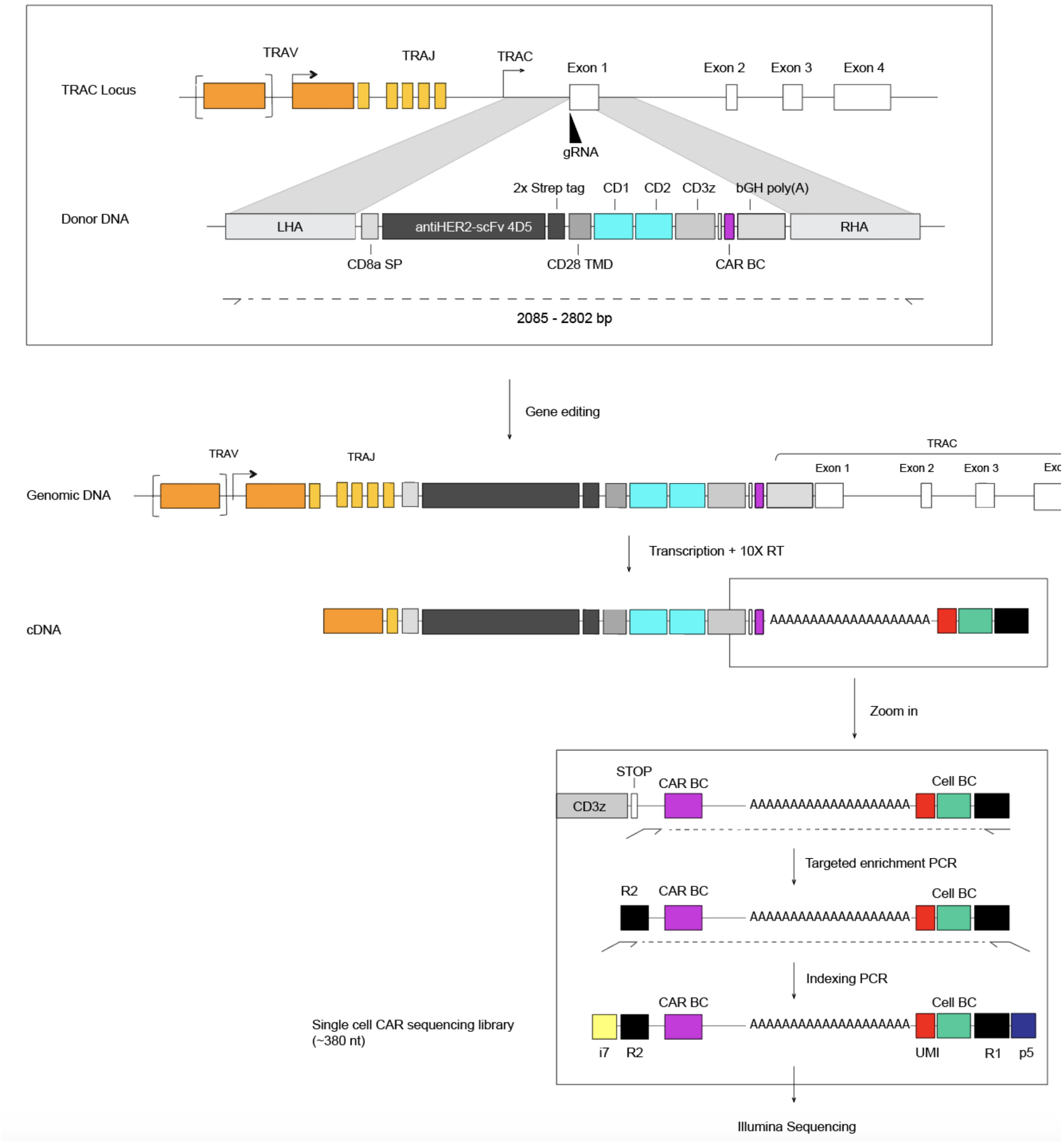

**Supp. Figure 3: scCARseq strategy for library demultiplexing**

Following CAR T cell engineering through CRISPR-Cas9 genome editing, a single copy of a CAR gene is integrated into the TRAC locus. In addition to the CAR components, each CAR gene contains a variant-specific barcode (purple) located in the 3' UTR region. The CAR gene is then transcribed into mRNA following *TRAC*-specific gene regulation. During the scRNAseq pipeline, mRNA is reverse transcribed to cDNA, and each transcript (including the CAR mRNA) incorporates a 3' barcode (green) specific to its cell of origin (cell-BC). In order to produce the scCARseq library, a two-step PCR amplification strategy first does a targeted amplification of the 3' UTR sequence of the CAR gene (that links the CAR-BC and the cell-BC) followed by an indexing PCR that adds the rest of the Illumina adaptors and sample index. The resulting library can then be sequenced using an Illumina platform to trace the origin of CAR transcripts and link them to the individual cells identified in the 10X gene expression pipeline.

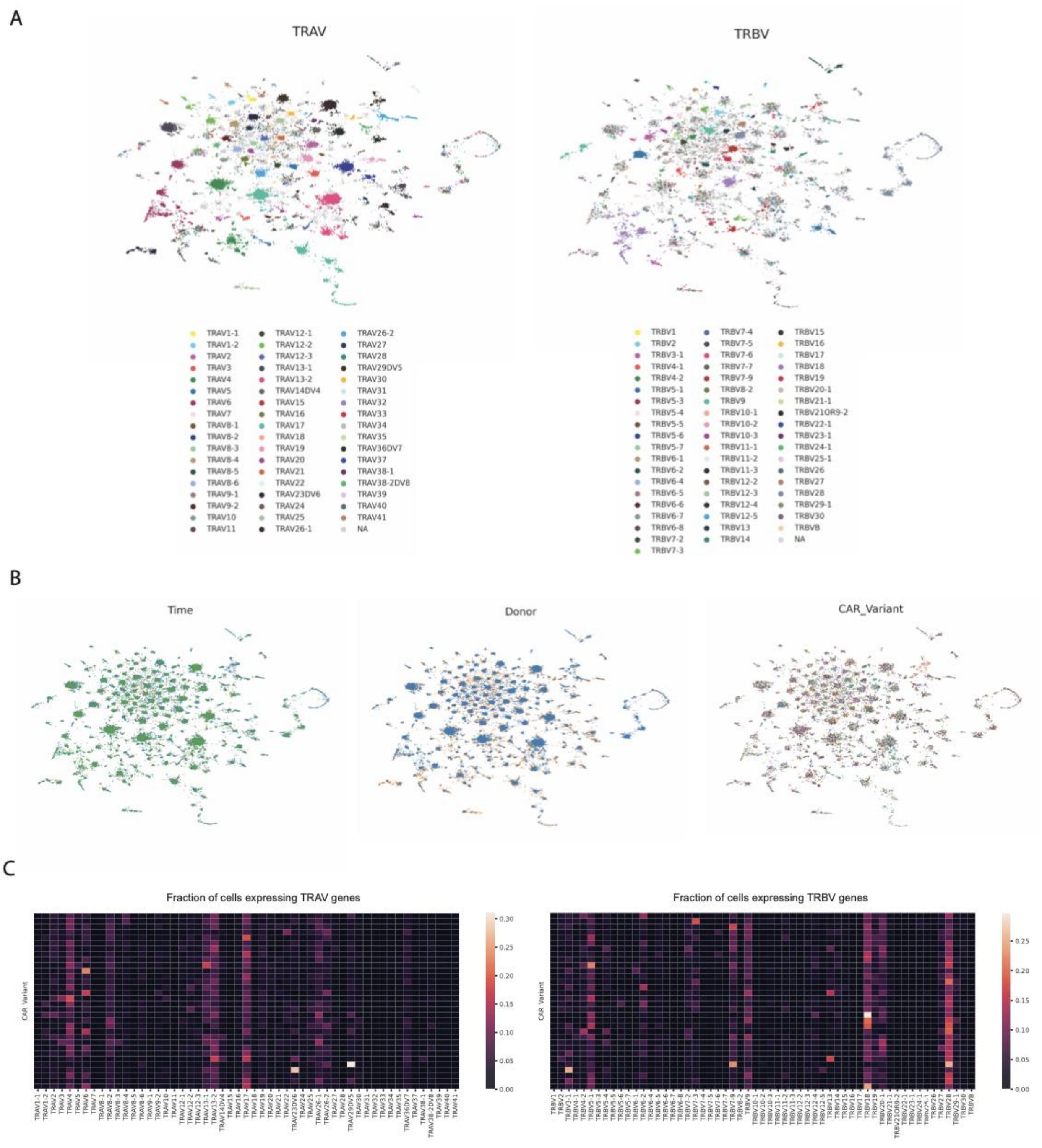

**Supp. Figure 4: Distribution of TCR variable genes across the CAR library scRNAseq data**

A- UMAP visualizations based on raw counts of TCR variable-alpha and variable-beta genes. Clustering in the UMAP space is driven by TRAV and TRBV germline gene segments (TRAJ and TRBJ included in UMAP learning but not shown).

B- UMAP visualization in A, coloured by time point, donor or CAR variant identity.

C- Heatmap illustrating the fraction of cells for each CAR variant, expressing each of the TRAV (left) or TRBV (right) genes.

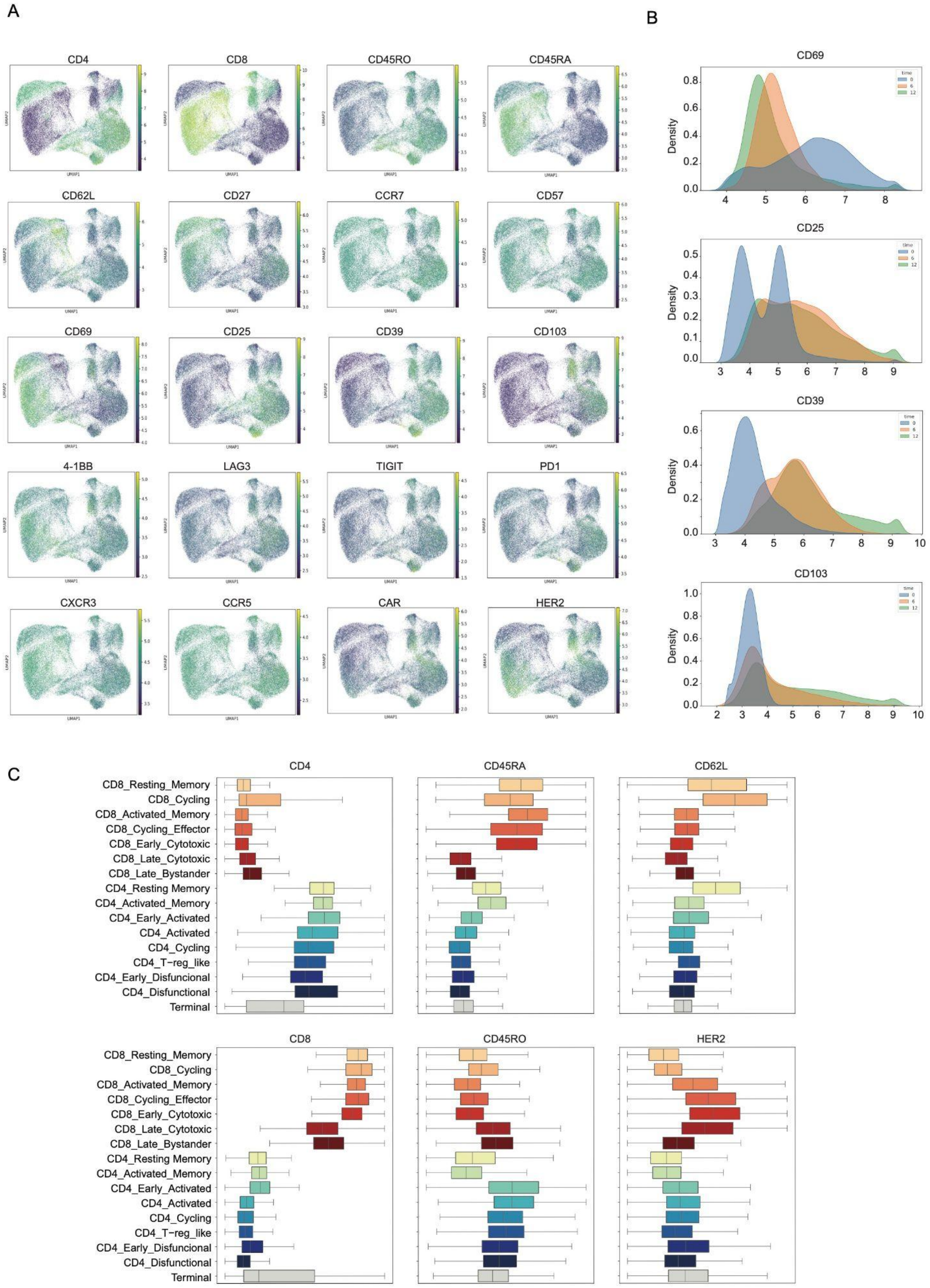
A
CD4
CD8
CD45RO
CD45RA
CD62L
CD27
CCR7
CD57
CD69
CD25
CD39
CD103
4-1BB
LAG3
TIGIT
PD1
CXCR3
CCR5
CAR
HER2
B
CD69
CD25
CD39
CD103
Density
time
0
6
12
C
CD4
CD45RA
CD62L
CD8
CD45RO
HER2
CD8_Resting_Memory
CD8_Cycling
CD8_Activated_Memory
CD8_Cycling_Effector
CD8_Early_Cytotoxic
CD8_Late_Cytotoxic
CD8_Late_Bystander
CD4_Resting Memory
CD4_Activated_Memory
CD4_Early_Activated
CD4_Activated
CD4_Cycling
CD4_T−reg_like
CD4_Early_Disfuncional
CD4_Disfunctional
Terminal

**Supp. Figure 5: scCITEseq characterization of T cell phenotypes**

**A**- UMAP embeddings from Figure 3B coloured based on the identification of different protein surface markers using scCITEseq. To increase contrast in the UMAPs, only dsb-normalized values between the 0.01 and 0.99 percentile are displayed, effectively removing the most extreme outliers that skew the color scale.

**B**- Change in dsb-normalized surface expression of early (CD69), middle (CD25) and late (CD39 and CD103) T cell activation markers across time.

**C**- Boxplot presenting the dsb-normalized surface display values for a selection of T cell marker genes across the different T cell clusters identified in Figure 3B.

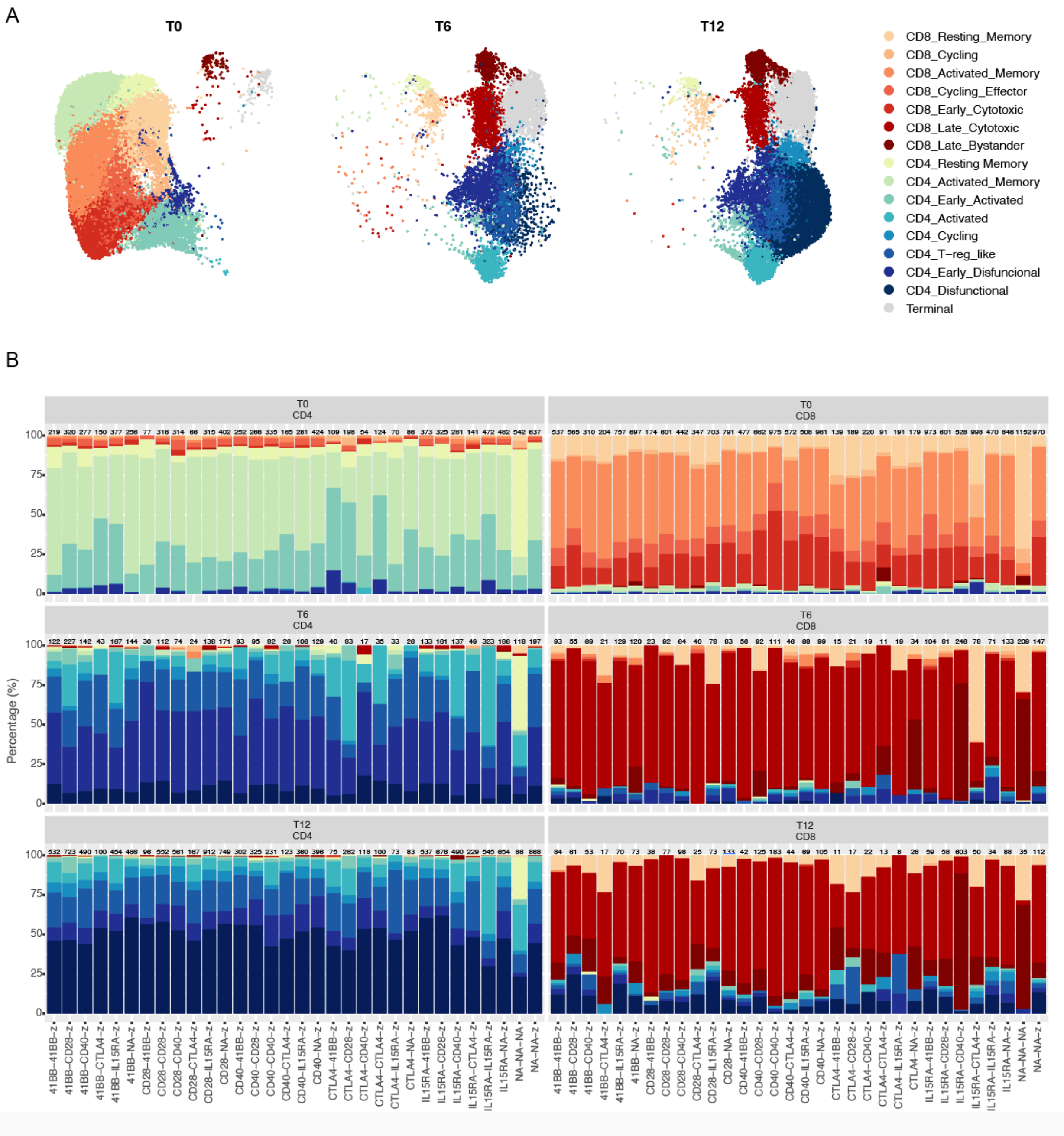

**Supp. Figure 6: Cluster enrichment across CAR variants, subsets and time**

**A**- UMAP embedding coloured by the clusters annotated in Figure 3 and split by the different time points of sample collection.
**B**- Cluster enrichment observed for the different CAR variants at early, middle and late time points of repeated tumor co-culture. The different time points for each CD8 or CD4 cell compartment are shown in different plots. The number of cells used to define the cluster distribution is reported at the top of each bar.

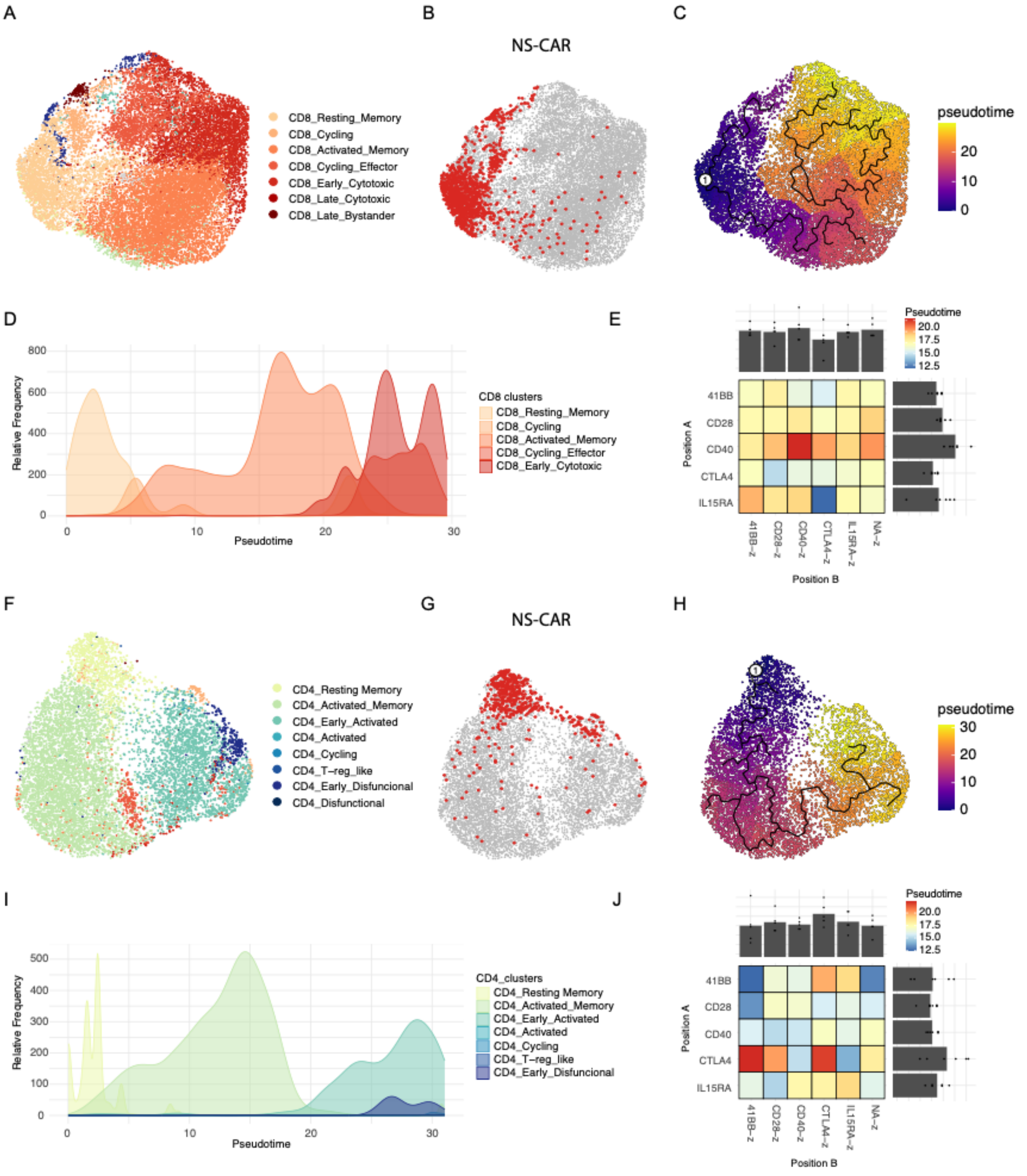

**Supp. Figure 7: Trajectories and pseudotime analysis of CAR T cell scRNAseq data at early activation orders cells by a T cell differentiation axis**

**A**- UMAP visualization of CD8 annotated cells at early time point, coloured by the cluster annotation from Figure 3.

**B**- UMAP embedding from (**B**) highlighting non-signaling CAR (NS CAR) annotated cells.

**C**- UMAP embedding from (**B**) showing a predicted cell trajectory depicted by a black line and coloured by predicted pseudotime. The trajectory was rooted manually by selecting the closest node to the NS-CAR population.

**D**- Distribution plot describing the cluster-specific relative frequency of cells along predicted pseudotime.
**E**- Heatmap showing the average pseudotime for CD8 CAR T cells during early activation across the different library variants. The heatmap separates variants based on the presence of CAR signaling domains in position A (proximal to the cell membrane) or position B (distal from cell membrane). In addition, bar plots at the top and right-hand side of the heatmap compile the pseudotime for all variants presenting a given domain in the different positions.
**F-J**- same as (**A-E)** but for CD4 cells.

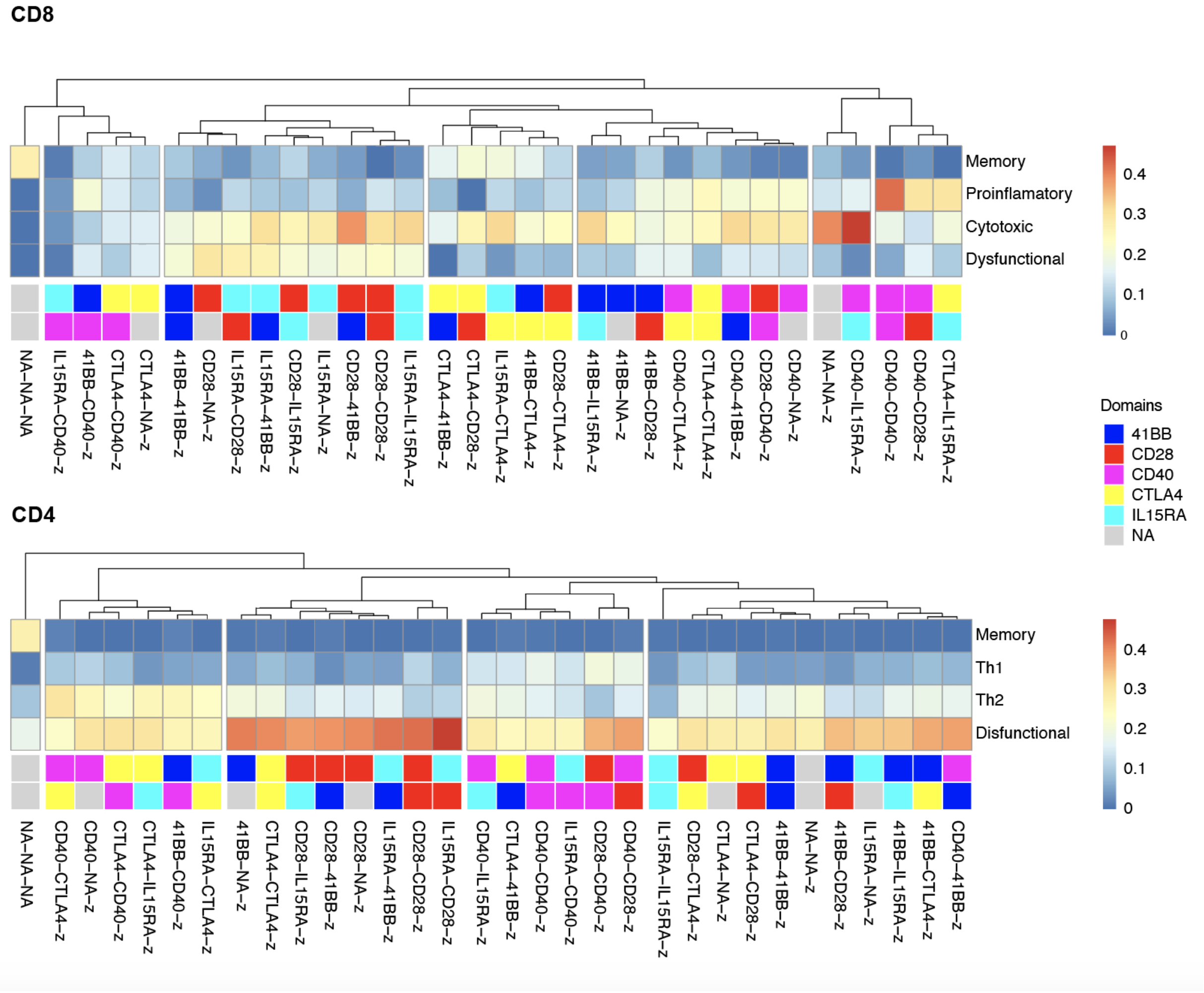

**Supp. Figure 8: Clustering of CAR variants according to functional clusters at late time point**

Hierarchical clustering of the CAR library according to the enrichment in late CD8 and CD4 clusters described in Figure 5 at the late time point (12 days) of the RAS assay.

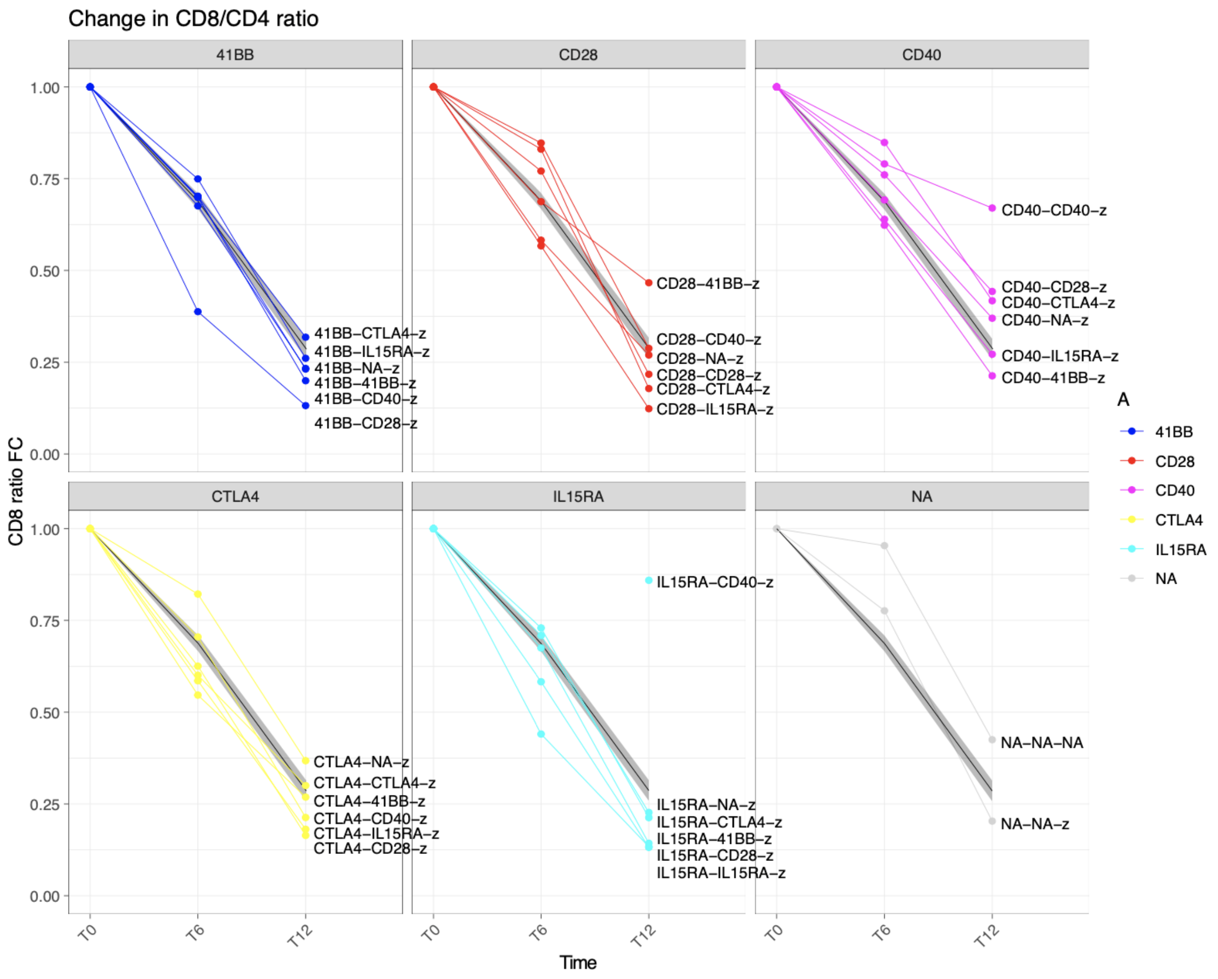

**Supp. Figure 9: CD8/CD4 ratio fold change over time across variants**

CD8 fold change (FC) throughout the progression of RAS based on scRNAseq data. Variants are coloured based on the domain located in a cell membrane-proximal position. A black line shows the mean FC with SEM in grey. FC was calculated by computing the difference in CD8/CD4 ratio compared to time-point 0.

SUPPLEMENTARY TABLES 1-5

**Supp. Table 1: CAR library nomenclature**

Comprehensive overview of the usage of intracellular signaling domains across the CAR library candidates. Positions A and B denote domains located proximal or distal to the cell membrane, respectively. NA indicates the absence of any domain in that given position. For each candidate, its associated molecular barcode is stated.

| Variant | Position A | Position B | CD3z | Barcode |
|---|---|---|---|---|
| 41BB-41BB-z | 41BB | 41BB | z | ACGGCGTTTCA |
| 41BB-CD28-z | 41BB | CD28 | z | GCCGACCTATA |
| 41BB-CD40-z | 41BB | CD40 | z | CCCGTCATCGG |
| 41BB-CTLA4-z | 41BB | CTLA4 | z | ATGGTCTTAAC |
| 41BB-IL15RA-z | 41BB | IL15RA | z | ATCGCACTGTA |
| 41BB-NA-z | 41BB | NA | z | GCCGGTCTAGT |
| CD28-41BB-z | CD28 | 41BB | z | CTCGGCCTAAC |
| CD28-CD28-z | CD28 | CD28 | z | AATGATATTAC |
| CD28-CD40-z | CD28 | CD40 | z | CCCGCAATCCG |
| CD28-CTLA4-z | CD28 | CTLA4 | z | AAGGAATTCTA |
| CD28-IL15RA-z | CD28 | IL15RA | z | GACGTTATAAA |
| CD28-NA-z | CD28 | NA | z | CCAGCTGTACT |
| CD40-41BB-z | CD40 | 41BB | z | CAAGTCCTGAG |
| CD40-CD28-z | CD40 | CD28 | z | GCAGGTCTGAC |
| CD40-CD40-z | CD40 | CD40 | z | GCCGCCATGCC |
| CD40-CTLA4-z | CD40 | CTLA4 | z | CCAGCCGTACA |
| CD40-IL15RA-z | CD40 | IL15RA | z | CGGGCAGTGCG |
| CD40-NA-z | CD40 | NA | z | TACGCTATTAA |
| CTLA4-41BB-z | CTLA4 | 41BB | z | AAGGATATTAG |
| CTLA4-CD28-z | CTLA4 | CD28 | z | GACGGTGTTAG |
| CTLA4-CD40-z | CTLA4 | CD40 | z | CCTGGCGTACG |
| CTLA4-CTLA4-z | CTLA4 | CTLA4 | z | AGTGGGGTTCA |
| CTLA4-IL15RA-z | CTLA4 | IL15RA | z | TCTGCGTTTCC |
| CTLA4-NA-z | CTLA4 | NA | z | GACGATATACG |
| IL15RA-41BB-z | IL15RA | 41BB | z | CTCGAAATGCA |
| IL15RA-CD28-z | IL15RA | CD28 | z | ATAGAAATCCC |
| IL15RA-CD40-z | IL15RA | CD40 | z | TAGGTAATGGA |
| IL15RA-CTLA4-z | IL15RA | CTLA4 | z | GCCGCGATCCA |

| Variant | Position A | Position B | CD3z | Barcode |
|---|---|---|---|---|
| IL15RA-IL15RA-z | IL15RA | IL15RA | z | ATCGAATTGTT |
| IL15RA-NA-z | IL15RA | NA | z | ACGGGTATACG |
| NA-NA-z | NA | NA | z | AAAGTGTTTAA |
| NA-NA-NA | NA | NA | NA | CCGGCACTATC |

**Supp. Table 2: Primers used in this study**

| Name | Sequence (5' to 3') | Purpose |
|---|---|---|
| F1 | GGTCAGACAAGCTCCCGGAAAAGGA | To amplify the cytoplasmic region of the CAR transgene integrated in the *TRAC*locus |
| R1 | AGGTGTCCCTTCCCTGCTT | |
| F2-mix | GTCACCTAAATGCTAGAGCTCGC | To amplify the 3' UTR region of the CAR transgene (containing a variant-specific barcode identifier). Primer degeneration is used to avoid Illumina sequencing problems due to sequence similarity. |
| | aGTCACCTAAATGCTAGAGCTCGC | |
| | tcGTCACCTAAATGCTAGAGCTCGC | |
| R2-mix | TACACGGCATGCCTGCTATTCT | |
| | aTACACGGCATGCCTGCTATTCT | |
| | tcTACACGGCATGCCTGCTATTCT | |
| F3 | GTGACTGGAGTTCAGACGTGTGCTCTTCCGATCTTGTCACCTAAATGCTAGAGCTCGCTG | scCAR-seq PCR1 |
| R3 | ACACTCTTTCCCTACACGACGCTC | |
| Dual Index Kit TT, Set A (PN-1000215) | AATGATACGGCGACCACCGAGATCTACAC-N10-ACACTCTTTCCCTACACGACGCTC | scCAR-seq PCR2 |
| | CAAGCAGAAGACGGCATACGAGAT-N10-GTGACTGGAGTTCAGACGTGT | |

**Supp. Table 3: Antibodies used in this study for flow cytometry**

| Target | Fluorochrome | Clone | Dilution | Source | Cat. Nr |
|---|---|---|---|---|---|
| StrepTag | Biotin | 5A9F9 | 1/200 | GenScript | A01737 |
| SAv | BV421 | - | 1/80 | Biolegend | 405226 |
| CD3ε | APC | UCHT1 | 1/200 | Biolegend | 300458 |

| | | | | | |
|---|---|---|---|---|---|
| Viability dye | Zombie NIR | - | 1/500 | Biolegend | 423105 |
| Viability dye | DRAQ7 | - | 1/200 | Biolegend | 424001 |
| CD4 | BV605 | OKT4 | 1/100 | Biolegend | 317437 |
| CD8a | BV711 | RPA-T8 | 1/100 | Biolegend | 301043 |
| CD107a | BV421 | H4A3 | 1/80 | Biolegend | 328626 |
| IFNγ | APC | B27 | 1/50 | Biolegend | 506510 |
| TNFα | PE | MAb11 | 1/200 | Biolegend | 502909 |

**Supp. Table 4: CITE-seq antibodies used in this study**

| Name | Target | Clone | Dilution | Source | Cat. Nr |
|---|---|---|---|---|---|
| TotalSeq™-B0953 PE Streptavidin | SAv | - | 1/400 | Biolegend | 405289 |
| TotalSeq™-B0072 anti-human CD4 | CD4 | RPA-T4 | 1/50 | Biolegend | 300565 |
| TotalSeq™-B0046 anti-human CD8 | CD8 | SK1 | 1/50 | Biolegend | 344757 |
| TotalSeq™-B0087 anti-human CD45RO | CD45RO | UCHL1 | 1/50 | Biolegend | 304257 |
| TotalSeq™-B0063 anti-human CD45RA | CD45RA | HI100 | 1/50 | Biolegend | 304161 |
| TotalSeq™-B0154 anti-human CD27 | CD27 | O323 | 1/50 | Biolegend | 302851 |
| TotalSeq™-B0148 anti-human CD197 (CCR7) | CCR7 | G043H7 | 1/50 | Biolegend | 353249 |
| TotalSeq™-B0085 anti-human CD25 | CD25 | BC96 | 1/50 | Biolegend | 302647 |
| TotalSeq™-B0168 anti-human CD57 | CD57 | QA17A04 | 1/50 | Biolegend | 393323 |
| TotalSeq™-B0146 anti-human CD69 | CD69 | FN50 | 1/50 | Biolegend | 310949 |

| | | | | | |
|---|---|---|---|---|---|
| TotalSeq™-B0088 anti-human CD279 (PD-1) | PD1 | EH12.2H7 | 1/50 | Biolegend | 329961 |
| TotalSeq™-B0176 anti-human CD39 | CD39 | A1 | 1/50 | Biolegend | 328241 |
| TotalSeq™-B0140 anti-human CD183 (CXCR3) | CXCR3 | G025H7 | 1/50 | Biolegend | 353751 |
| TotalSeq™-B0141 anti-human CD195 (CCR5) Antibody | CCR5 | J418F1 | 1/50 | Biolegend | 359139 |
| TotalSeq™-B0355 anti-human CD137 (4-1BB) | 41BB | 4B4-1 | 1/50 | Biolegend | 309837 |
| TotalSeq™-B0133 anti-human CD340 (erbB2/HER-2) | HER2 | 24D2 | 1/50 | Biolegend | 324425 |
| TotalSeq™-B0145 anti-human CD103 (Integrin αE) | CD103 | Ber-ACT8 | 1/50 | Biolegend | 350235 |
| TotalSeq™-B0152 anti-human CD223 (LAG-3) | LAG3 | 11C3C65 | 1/50 | Biolegend | 369337 |
| TotalSeq™-B0089 anti-human TIGIT (VSTM3) | TIGIT | A15153G | 1/50 | Biolegend | 372727 |
| TotalSeq™-B0147 anti-human CD62L | CD62L | DREG-56 | 1/50 | Biolegend | 304849 |

**Supp. Table 5: Gene sets used for gene set scoring**

| **Cytotoxicity** | **Proinflamatory** | **Memory** | **CD4_Th1** | **CD4_Th2** |
|---|---|---|---|---|
| GZMB | IFNG | TCF7 | IL2 | IL5 |
| PRF1 | TNF | SELL | IFNG | IL13 |
| FASLG | CRTAM | CCR7 | TNF | IL4 |
| | CSF2 | LEF1 | | |
| | XCL1 | IL7R | | |
| | XCL2 | | | |

| | |
|---|---|
| | CCL1 |
| | CCL4 |
